# Tenascin-C in the early lung cancer tumor microenvironment promotes progression through integrin αvβ1 and FAK

**DOI:** 10.1101/2024.09.17.613509

**Authors:** Shiela C. Samson, Anthony Rojas, Rebecca G. Zitnay, Keith R. Carney, Wakeiyo Hettinga, Mary C. Schaelling, Delphine Sicard, Wei Zhang, Melissa Gilbert-Ross, Grace K. Dy, Michael J. Cavnar, Muhammad Furqan, Robert F. Browning, Abdul R. Naqash, Bryan P. Schneider, Ahmad Tarhini, Daniel J. Tschumperlin, Alessandro Venosa, Adam I. Marcus, Lyska L. Emerson, Benjamin T. Spike, Beatrice S. Knudsen, Michelle C. Mendoza

## Abstract

Pre-cancerous lung lesions are commonly initiated by activating mutations in the RAS pathway, but do not transition to lung adenocarcinomas (LUAD) without additional oncogenic signals. Here, we show that expression of the extracellular matrix protein Tenascin-C (TNC) is increased in and promotes the earliest stages of LUAD development in oncogenic KRAS-driven lung cancer mouse models and in human LUAD. TNC is initially expressed by fibroblasts and its expression extends to tumor cells as the tumor becomes invasive. Genetic deletion of TNC in the mouse models reduces early tumor burden and high-grade pathology and diminishes tumor cell proliferation, invasion, and focal adhesion kinase (FAK) activity. TNC stimulates cultured LUAD tumor cell proliferation and migration through engagement of αv-containing integrins and subsequent FAK activation. Intringuingly, lung injury causes sustained TNC accumulation in mouse lungs, suggesting injury can induce additional TNC signaling for early tumor cell transition to invasive LUAD. Biospecimens from patients with stage I/II LUAD show TNC in regions of FAK activation and an association of TNC with tumor recurrence after primary tumor resection. These results suggest that exogenous insults that elevate TNC in the lung parenchyma interact with tumor-initiating mutations to drive early LUAD progression and local recurrence.

## Introduction

Lung cancer is the deadliest cancer worldwide^1^. Lung adenocarcinoma (LUAD) is the most prevalent form of lung cancer. While stage 0 adenocarcinoma *in situ* and minimally invasive adenocarcinoma have a 98% survival rate after resection^2^, nearly one third of the more invasive stage I/II lung adenocarcinomas (LUADs) recur^3^. As a result, stage I and II lung cancer patients face 5 year survival rates of 85.6% and 66.5%, respectively^4^. Once disseminated (stage III/IV), survival drops under 40% and 10% respectively^4^. Early lung cancers can lie dormant for several decades^5^. Screening efforts aim to reduce lung cancer mortality by catching and surgically resecting cancers before they have spread throughout the body^6^. A better understanding of the processes that convert indolent lesions to aggressive LUAD could identify novel prognostic biomarkers and therapeutic targets that guide treatment decisions and reduce the risk of recurrence.

Activating mutations in upstream components of the RAS→RAF→MEK→ERK pathway^7^ are the earliest events that intiate most LUAD. These initial mutations can be present for decades before cancer develops^5, 8, 9^. Additional genetic hits, most commonly the loss of tumor suppressors *TP53* or *LKB1*, increase in frequency during clinical progression in patients, and drive malignant progression to early cancer and metastasis in mouse models^7, 10^. *LKB1* mutations are less common than *TP53* mutations, but result in more rapidly aggressive tumors in genetically-engineered mouse models (GEMMs)^11–13^. The aggressiveness of *LKB1*-mutant LUAD is attributed, in part, to activation of focal adhesion kinase (FAK), which promotes invasion and collagen remodeling^13, 14^. Yet, the time needed for cancer to develop in these mouse models and heterogeneity in the process suggest that epigenetic or environmental changes also contribute to early oncogenesis.

Adaptive oncogenesis posits that pre-cancerous cells with initiating mutations lie dormant until awakened by ageing or environment-induced changes in the host tissue^15, 16^. Through inhalation, the lung is in regularly exposed to environmental irritants and toxins that can cause injury. Indeed, lung cancer incidence increases with age, smoking, exposure to air pollution, fibrosis, and radiation therapy near the lung with associated lung scarring^1, 17–21^. These risks are associated with non-cell autonomous changes in the tumor microenvironment: inflammatory stress, reduced immune surveillance, and increased extracellular matrix (ECM) deposition and crosslinking^8, 22–25^. Among the ECM components, the glycoprotein Tenascin-C (TNC) is noteable for its low, rare expression in adult lung and dramatically increased expression during lung injury. TNC expression normally occurs during fetal and newborn lung development and is lost by early adulthood^26–28^. Exposure to bleomycin, which damages the lung, induces acute TNC re-expression^22, 25, 29, 30^. TNC expression has also been observed in an LUAD patient after radiation therapy^31^. This raises the intriguing possibility that throughout one’s lifetime, lung damage may repeatedly cause TNC expression, which could affect cancer development and recurrence.

In models of metastatic breast cancer, TNC expression induced stemness and metastatic outgrowth in the lung^32, 33^. In a glioma model, TNC induced tumor stiffness, mechanosignaling with FAK activation, and aggressiveness^34^. In LUAD transplant mouse models, *TNC* expression drives the seeding and metastasis of advanced LUAD cells into the lung^30^. In LUAD patients, *TNC* expression correlates with poor survival^30, 35^. The contribution of TNC to the transition and early progression of benign adenomas to lung cancer remains unknown.

Here, we sought to determine how TNC contributes to the transition of early precancerous adenomas to LUAD. Using KRAS-driven GEMMs of LUAD, human clinical samples, and experiments with purified TNC, we found that TNC expressed by adenoma- and transitioning LUAD-associated fibroblasts activated integrin αvβ1 and FAK in tumor cells to drive tumor progression.

## Results

### TNC is expressed by fibroblasts in the early LUAD tumor microenvironment

To better understand the role of TNC in early LUAD development, we examined TNC expression in early tumors using GEMMs of RAS-driven LUAD. These models allowed us to capture the earliest stages of tumor development that occur before symptoms develop and before most clinical diagnoses. In *KRas^LSL-G12D/+^; Rosa26^LSL-YFP^* (*KY*) and *KRas^LSL-G12D/+^; Rosa26^LSL-tdTomato^* (*KT*) mice, intratracheal intubation of Cre adenovirus induces somatic expression of oncogenic KRAS^G12D^ along with expression of a fluorescent protein label. KRAS^G12D^ expression initiates multi-focal tumor development, with atypical adenomatous hyperplasia (AAH) presenting after 2-5 weeks and early adenomas presenting after 12 weeks^10, 36, 37^. In mice that additionally lose TP53 upon Cre administration (*KRas^LSL-G12D/+^;Trp53^F/F^;Rosa26^LSL-tdTomato^*, *KPT*), AAH and adenomas begin to transition to LUAD at 10-12 weeks^10, 36, 37^ (Fig. 1a and Supplementary Fig. 1a, b). The LUADs invade the lung parenchyma and become metastatic by ∼26 weeks^9,11,3^. Histopathological analysis and sequencing have shown that the GEMM tumor progression generally replicates clinical LUAD, in which pre-invasive stage 0 adenomas progress to stage I/II LUADs with heterogenous histology, dedifferentiation, and local invasion^10, 37, 38^.

**Figure 1.**
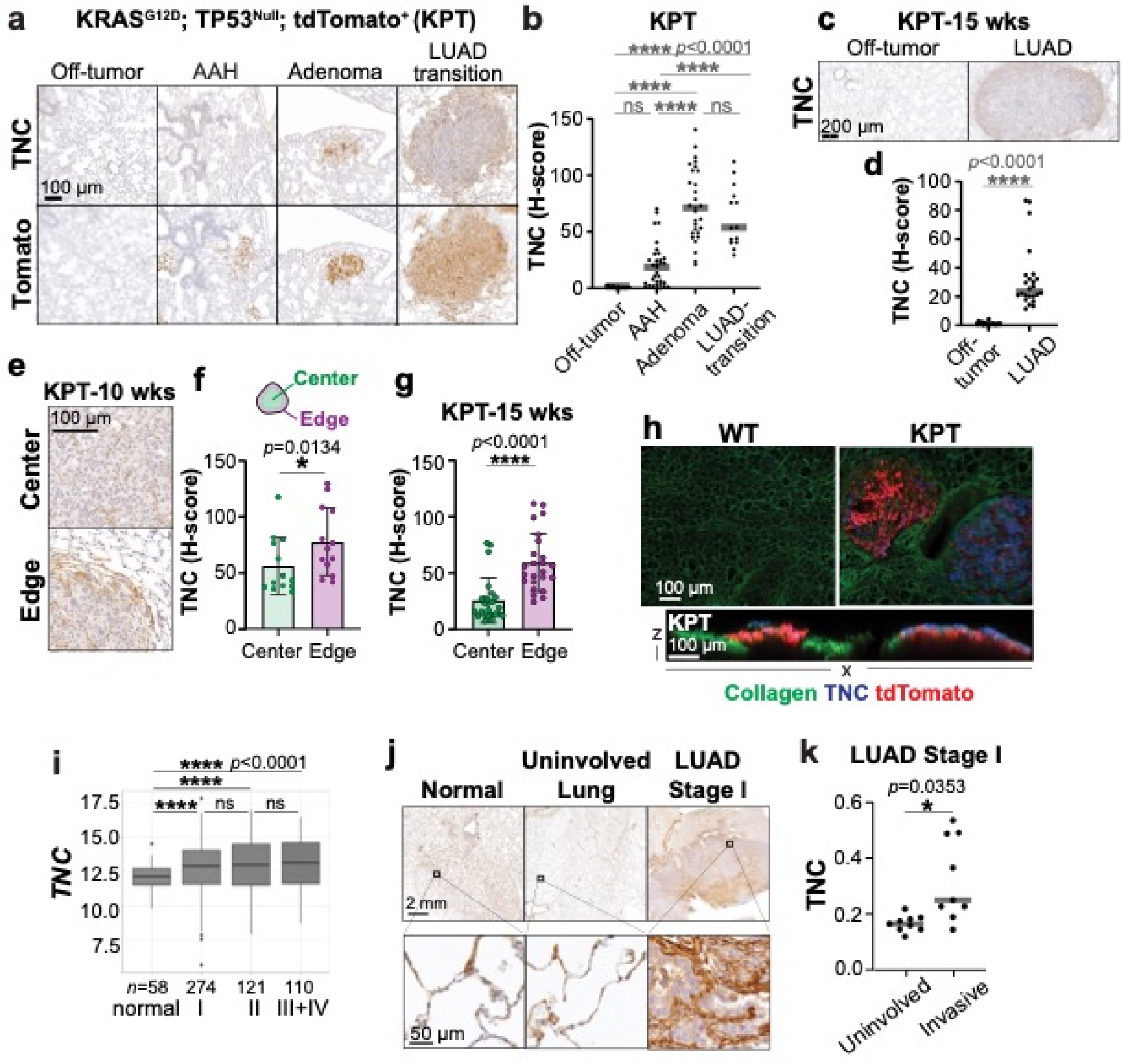
TNC is expressed at the tumor-stroma interface in early LUAD. **a** Representative lung lesions from *KPT* mice 10 weeks after Cre inoculation. Immunohistochemistry on serial sections for transformed cells (tdTomato+) and TNC, *n*=3 females and 3 males. **b** Quantification of TNC in (a). Points are TNC intensity within individual lesions. One-way ANOVA with Tukey’s posthoc test. **c** Representative immunohistochemistry for TNC in lung lesions from *KPT* mice 15 weeks after Cre inoculation, *n*=4 females and 4 males. **d** Quantification of TNC in (c). One-way ANOVA with Tukey’s posthoc test. **e** Representative immunohistochemistry for TNC at tumor center and edge in transitioning LUADs in (a). **f, g** Quantification of TNC at tumor center and edge for 10 week transitioning LUADs and 15 week established LUADs from *KPT* mice in (b, d). Kruskall-Wallis test. **h** Representative immunofluorescence for TNC and collagen labelling with CNA35-GFP in PCLS’s from KPT LUAD, 15 weeks, *n*=3 mice. KPT tumor on left shows low TNC and tumor on right shows high TNC. **i** *TNC* mRNA in LUAD, as a function of tumor stage, from TCGA. Counts are normalized Log2 fold change. **j** Representative immunohistochemistry for TNC in normal human lung tissue (autopsy) and uninvolved lung and tumor tissue from LUAD resections. Uninvolved lung is >4 cm from tumor boundaries and histologically benign, *n*=5 autopsy, *n*=4 uninvolved, and *n=*11 stage I LUAD. **k** Intensity of stromal TNC from human lung resections with both uninvolved lung and invasive LUAD regions, *n*=9. Points are mean intensity per case. One-way ANOVA.

We performed immunohistochemistry for TNC on lung tissue from the *KPT* model at 10 weeks, when the transition from adenoma to LUAD becomes microscopically apparent, and at 15 weeks, when both adenomas transitioning to LUAD and established, invasive LUADs are present (Supplementary Fig. S1a). TNC expression was increased in the earliest adenoma and transitioning LUAD stages at 10 weeks (Fig. 1a, b). TNC remained elevated in the established LUADs present at 15 weeks (Fig. 1c, d). The off-tumor tissue of KPT mice exhibited normal alveolar structure and no detectable TNC expression, similar to lungs from control mice with wildtype (WT) *Kras* (*Kras^+/+^;Trp53^F/F^; Rosa^LSL-tdTomato^*) (Supplementary Fig. 1c). Since fibrotic collagen deposition following lung injury is more severe in male mice than in female mice^39^, we compared TNC expression in LUAD by sex. Interestingly, TNC expression was also greater in the early LUADs from male mice compared to those in female mice (Supplementary Fig. 1d).

We noted that TNC was expressed primarily at the tumor edge rather than the tumor center (Fig. 1a, c, e). Quantification of TNC staining at the edge versus the center of transitioning LUADs at 10 weeks and established LUADs at 15 weeks showed TNC expression to be primarily at the tumor edge, where the tumor interfaces with the stroma (Fig. 1f, g). Staining for TNC in precision-cut lung slices (PCLS) of the established LUADs and 3D confocal scanning confirmed that TNC was primarily located outside the circumference of tdTomato+ tumor masses, regardless of whether TNC expression was relatively low or high (Figure 1h). This suggested that TNC is likely produced in the tumor microenvironment, rather than by the tumor cells themselves, at this early stage of LUAD. We tested if TNC is also increased at the edges of tumors generated in the less agressive *KT* model and a more aggressive model, driven by activation of KRAS^G12D^ and loss of LKB1 (*Kras^LSL-G12D/+^; Lkb^FlF^* (*KL*) mice)^13^. TNC was present at the edge of KT adenomas, although the intensity was lower than in KPT tumors (Supplementary Fig. 1e, f). TNC was also present in transitioning KL LUADs and in this case, the staining was again stronger in tumors from male mice than female mice (Supplementary Fig. 1g-i).

We next tested if TNC is expressed in the early stages of human LUAD. RNAseq data from The Cancer Genome Atlas (TCGA)^40^ showed *TNC* mRNA was increased in stage I samples, compared to normal lung (Fig. 1i). *TNC* expression remained high in later stage tumors (Fig. 1i). We confirmed this pattern by immunohistochemistry on clinical samples obtained from the University of Utah Department of Pathology. We assayed histologically benign, or normal, lung tissue from unrelated autopsies as well as benign, uninvolved and tumor tissue from surgical resections from lung cancer patients. In most cases, histologically benign lung tissue showed weak TNC expression, while stage I LUAD exhibited increased TNC (Fig. 1j). Grouping the cases into categories of weak, moderate, and strong TNC expression revealed that ∼¾ of benign lung tissue exhibited weak expression, while nearly all stage I, II, and III/IV cases exhibited moderate or high TNC expression (Supplementary Fig. 1j, k). We systematically compared the TNC intensity in invasive regions of small tumors (T1/T2) to histologically normal off-tumor regions of the same samples. In order to specifically examine regions of tumor-stroma interface, we applied machine learning to automatically identify and exclude tumor and immune cell areas from the TNC intensity calculations (Supplementary Fig. 1l). The analysis showed that invasive regions of LUAD had significantly more TNC than non-tumor regions within the same tissue section (Fig. 1k).

We utilized the KPT model to probe the source and signaling of TNC in LUAD, since *Trp53*-mutant LUADs are more common than *LKB1*-mutant LUADs in clinical patients and their less aggressive progression when modeled with KRAS mutations results in a larger number of early and transitional lesions^11^. A previous study showed that advanced KP tumors cells express TNC^30^. However, quantification of *TnC* mRNA from ssRNA-sequencing of KP LUADs 16 weeks after initiation^41^ showed that fibroblasts expressed significantly more *TnC* than other cell types, including tumor and immune cells (Supplementary Fig. 2a). We used *in situ* hybridization (ISH) to detect *TnC* mRNA production in the KPT tumors and immunohistochemistry co-detection for Tomato to identify the tumor cells. Immunohistochemistry for vimentin and elongated cell morphology identified the fibroblasts. *TnC* signal was interspersed between tdTomato-positive tumor cells and overlapped with elongated, tdTomato-negative, vimentin-positive cells in early LUAD (Fig. 2a. b). The lack of *TnC* signal in tdTomato positive cells suggests a stromal source of *TnC* rather than cancer cells that have undergone epithelial-to-mesenchymal transition (EMT). Since vimentin is also expressed by other cell types in the mesenchymal lineage, we queried the integrated Human Lung Cell Atlas (HLCA) for TNC^42^ to identify additional lung fibroblast markers that identify the *TNC+* cell population. Of the vimentin+ cell types in normal lung tissue, a small subset of alveolar type 1 fibroblasts expressed the highest levels of *TNC* and a subset of myofibroblasts express a lower level of *TNC* (Supplementary Fig. 2b, c). Alveolar fibroblasts can transition to “activated fibroblasts” that express myofibroblast-like genes, including *Acta2* (α *Smooth muscle actin*, *αSMA)*, in response to lung injury^22, 25, 42^. We found that αSMA was expressed in regions at the KPT early tumor edge that overlap with TNC and vimentin (Fig. 2c, d). This suggests that in early LUAD, TNC expression is mediated by a rare population of activated alveolar type 1 fibroblasts or myofibroblasts, not other vimentin+ mesenchymal cells.

**Figure 2.**
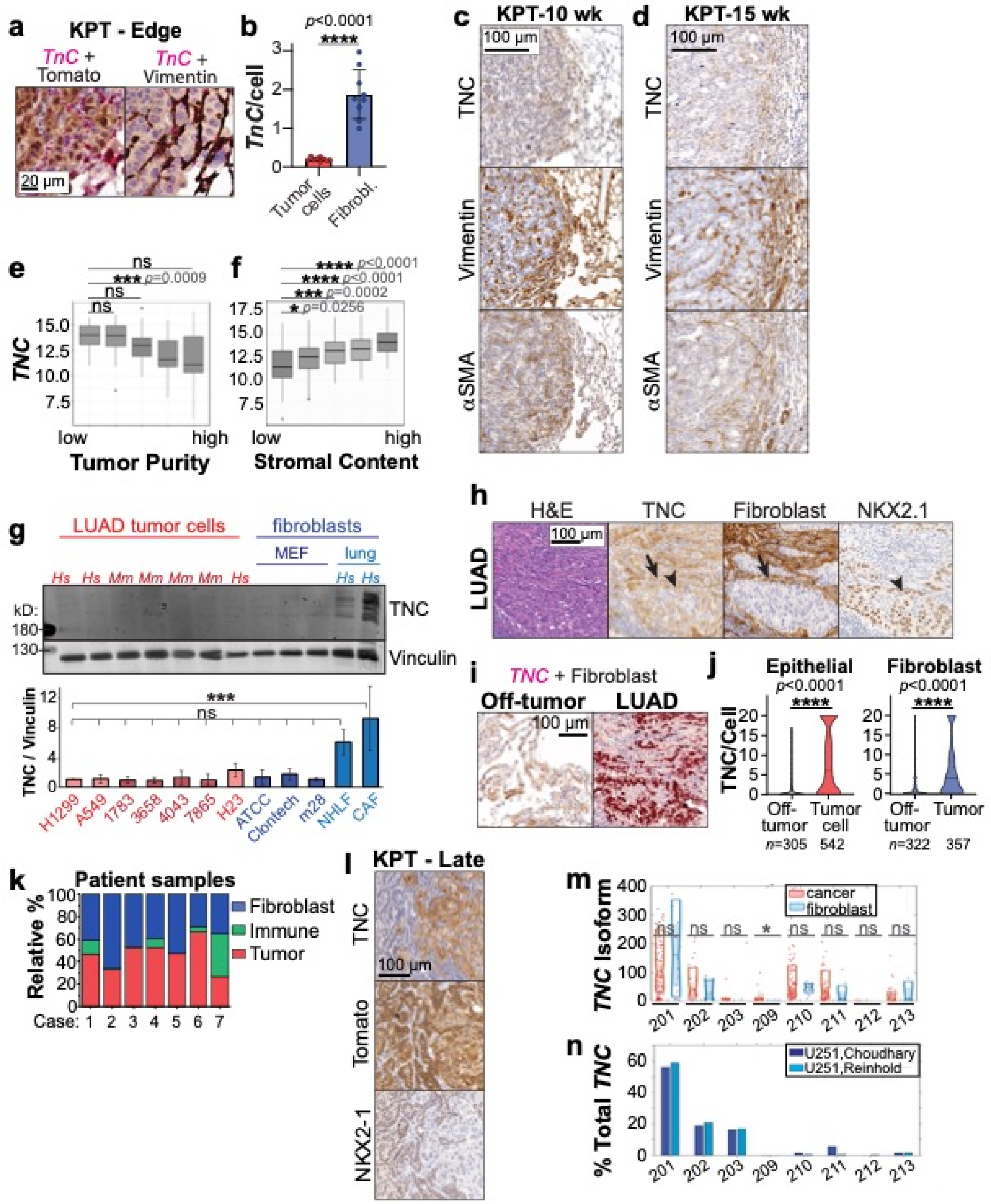
Fibroblasts at the tumor-stroma interface express TNC. **a** Representative KPT LUAD, 10 weeks, with *TnC* ISH (red) and immunohistochemistry co-detection of tdTomato or Vimentin (DAB-brown). 9 ROIs, from 8 tumors in *n*=3 mice, female. **b** Quantification of *TnC* particles in (a). Two-tailed T-test with Welch’s correction. **c, d** Representative immunohistochemistry for TNC and fibroblast markers in serial sections of transitioning LUAD, *n*=3 female and 3 male KPT mice at each time point. **e, f** *TNC* mRNA in human LUAD, as a function of tumor purity and stromal content, TCGA Pan Cancer Atlas. Counts are normalized Log2 fold change. **g** Representative western blot and quantification of TNC in LUAD tumor cell lines and fibroblasts, *n*=4. Human LUAD cell lines H1299, A549, H23. Mouse LUAD cell lines 1783, 3658, 4043, 7865. MEF, mouse embryonic fibroblast. NHLF, normal human lung fibroblast, immortalized. CAF, lung cancer-associated fibroblasts. Error bars SD. **h** H&E and immunohistochemistry images from human stage I LUAD. Off-tumor is tumor >1 mm from tumor boundary. Representative of *n*=15 cases of stage I or II LUAD. Arrow shows TNC overlapping with fibroblast regions and arrowhead TNC overlapping with tumor cells bordering fibroblasts. **i** Representative ISH-immunohistochemistry codetection of *TNC* (red) and fibroblasts (TE7 antibody, DAB-brown), *n*=7 cases of stage I and II LUAD. **j** Distribution of *TNC* particles/cell in off-tumor and LUAD ROIs in (i), manual counting. K-S test. *n=*cells. **k** Distribution of *TNC* expression in tumor, stroma and immune regions for cases in (i), automated detection described in Supplementary Fig. 2f). **l** Representative immunohistochemistry for TNC and tumor cell markers tdTomato and NKX2-1 in serial sections of late KPT tumors. **m** *TNC* isoforms in LUAD. RNAseq by Expectation Maximization (RSEM) expected counts from RNAseq of primary LUAD patient samples, pre-treatment^48^. **n** *TNC* isoforms expressed in U251 cells. RSEM, calculated from RNAseq of U251 cells^49, 50^ and plotted as a % total *TNC*.

We tested if fibroblasts also drive TNC expression in the early-presenting human lung cancer. ESTIMATE scores for tumor purity and stromal content^43^ showed that human LUAD samples with high stromal content and low tumor purity expressed more *TNC* than samples with low stromal content and high tumor purity (Fig. 2e, f), suggesting fibroblasts express TNC in human LUAD. Western blotting for TNC across a panel of mouse and human LUAD tumor cells and fibroblasts showed that *in vitro*, LUAD tumor cells express nearly undetectable TNC, while lung cancer-associated fibroblasts (CAFs) express significant levels of TNC (Fig. 2g). We then stained serial sections of our clinical LUAD samples for TNC, the lung lineage factor NKX2-1 to label early tumor cells^44^, and TE-7 to label fibroblasts^45^. TE-7 labels fibroblasts and myofibroblasts without also staining CD31+ endothelial cells (Supplementary Fig. 2d). In invasive regions at the edge of stage I and II tumors, TNC expression overlapped with fibroblasts and also NKX2-1+ tumor cells bordering the fibroblast/TNC region (Fig. 2h). ISH with immunohistochemistry co-detection of fibroblasts and manual counting of *TNC* particles showed that human clinical samples, both tumor cells and fibroblasts expressed significantly more *TNC* in invasive regions of LUAD than in off-tumor regions (Fig. 2i, j). We also applied automated quantification of *TNC* mRNA in regions of interest (ROIs) stochastically-selected from benign lung and the invasive edge of LUAD in the serial section H&E stains. We generated masks of TE7+ fibroblasts and tumor and immune cells and quantified the number of *TNC* mRNAs objects within segmented cells within each ROI (Supplementary Fig. 2f). In seven cases of invasive stage I or II LUAD, both fibroblasts and tumor cells produced *TNC* (Fig. 2k). We noted that TNC-positive tumor cells retained expression of the lung lineage factor NKX2-1+ and were adjacent to extracellular TNC fibers (Fig. 2h, i).

That fibroblasts and NKX2-1+ tumor cells can both express TNC in early LUADs differs from a previous report of high-grade LUADs, which describes tumors cells as the primary source of TNC^30^. The reports shows that tumor cells can express TNC as a result of loss of NKX2-1^30^, contrary to our observation in early stage mouse model and human LUAD (Fig. 2a, b, h-j). To determine the association of NKX2-1 and TNC expression, we tested TNC expression in more advanced tumors from our mouse models. We confirmed that in mice 26 weeks after tumor induction, TNC staining overlaps with NKX2-1-negative tumor cells (Fig. 2l). Deletion of *Nkx2-1* in the lung epithelial cells at the start of tumor initiation in *KRas^LSL-G12D/+^;Nkx2-1^F/F^; Rosa^LSL-tdTomato^* (KNT) mice^46^ resulted in low-level TNC expression throughout early tumors and increased expression in late tumors, confirming that NKX2-1 suppresses *TNC* expression (Supplementary Fig. 2e). Single cell-sequencing of tumors from the KPT model has shown that *Nkx2-1*-silenced tumor cells arise with the development of invasive LUAD and expand in high-grade LUAD^10^. While the translation of these data to human LUAD is imperfect in that human stage I LUAD is more advanced than the transitioning LUAD assayed in mice^47^, we conclude that TNC is initially expressed by lung fibroblasts at the edge of early tumors and that early tumor cells can be locally activated to express TNC and can gain further TNC expression as NKX2-1 expression decreases. Since TNC is expressed as multiple isoforms and larger splice variants are preferentially expressed in other solid tumors, we probed published scRNAseq data from human LUAD^48^ to identify the *TNC* isoforms produced by LUAD tumors cells versus fibroblasts. The *TNC* isoforms were expressed at nearly identical levels in tumor cells and fibroblasts, except for low versus no expression of isoform 209 (Fig. 2m). The predominant isoform was 201, which is the same isoform commercially isolated from glioblastoma U251 cells^49, 50^ (Fig. 2n). Thus, TNC’s function in early LUAD is unlikely to be impacted by whether it is produced by activated fibroblasts or tumor cells.

### TNC induces tumor cell aggressiveness by activating integrin αvβ1 and FAK

We tested if TNC expression is functionally important in early LUAD. Gene set enrichment analysis (GSEA) using the TNC-response gene set on TCGA data showed that LUADs with the most *TNC* mRNA expression exhibited increased expression of TNC-response genes^51^ (Supplementary Fig. 3a). The core group of genes that accounted for the enchrichment clustered into gene ontology (GO) terms for biological processes of cell signaling, cell adhesion, and cell migration (Supplementary Tables 1 and 2). Consistent with these enrichments, human LUAD tumor cells (H1299) cultured on TNC-coated plates exhibited increased cell proliferation and migration (Fig. 3a-c).

**Figure 3.**
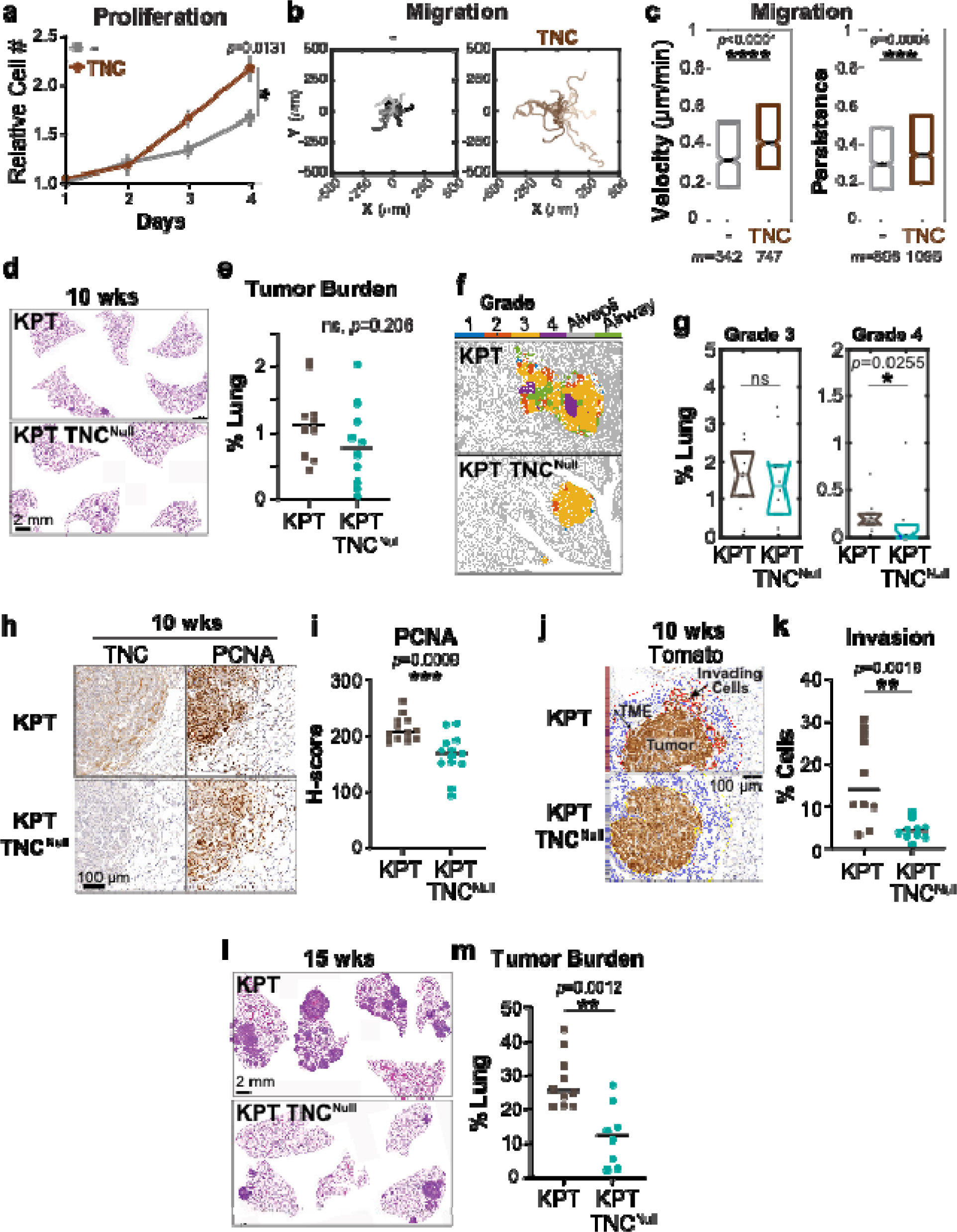
TNC promotes early LUAD progression. **a** H1299 cell proliferation on plates coated with bovine serum albumin (BSA) or TNC. Error bars are SEM for *n*=6 independent experiments. One-way ANOVA with Tukey’s test. **b, c** H1299 cell migration on plates coated with BSA or TNC. Rose plots show tracks of 20 representative cells. Box plots are *m* cells from *n*=3 independent experiments. K-S test. **d** Representative H&E image of KPT and KPT TNC^Null^ lung lesions, 10 weeks after Cre inoculation, *n*=5 female, 5 male KPT and 5 female, 5 male KPT TNC^Null^. **e** Quantification of tumor burden for (d). **f, g** Representative image of GLASS-AI grade segmentation and quantification for (d). Normal alveoli and airways are classified in grey and green, respectively. **h, i** Representative immunohistochemistry for PCNA and quantification at the tumor edge for samples in (d). **j** Representative identification of tumor cells in tumor microenvironment region (cell borders outlined in red) outside of primary tumor using tdTomato immunohistochemistry. **k** Quantification of percentage of invading tumor cells in (j). **l** Representative H&E on serial sections of lung lesions from KPT and KPT TNC^Null^ mice 15 weeks after Cre inoculation. *n*=5 female, 5 male KPT lungs and 4 female, 4 male KPT TNC^Null^. **m** Quantification of tumor burden in (j). Tumor burden, grade, PCNA, and invasion tested with unpaired T-test.

Next, we tested if TNC in the host environment affects early primary tumor growth *in vivo*. First, we orthotopically transplanted syngeneic, mouse LUAD tumor cells (3658) derived from high-grade KNT tumors^46^ into the lungs of *TnC* wildtype (WT) and *TnC* knockout (KO) mice. 3658 cells express low levels of TNC, compared to human and mouse lung fibroblasts (Fig. 2g). The TNC^Null^ lungs of adult *TnC* KO mice are grossly and functionally normal^52, 53^, but harbor less αSMA expression and increased TGFβ signaling^54^. 4 weeks after transplantation, tumor burden in TNC^Null^ lungs was significantly lower than in WT lungs (Supplementay Fig. 3b-d). In tumors from the TNC WT mice, TNC was expressed at the tumor-stroma interface, in regions that overlapped with vimentin+ fibroblasts. In the tumors grown in TNC^Null^ lungs, weak TNC expression occurred within the tumors, due to the weak expression of TNC by 3658 cells (Supplementay Fig. 3c). We then crossed the KPT mice with *TnC* KO mice and induced tumor growth to generate KPT TNC^Null^ tumors. At 10 weeks, the overall tumor burden in the KPT-TNC^Null^ lungs trended lower than that of KPT lungs, but was not significantly different (Fig. 3d, e). The KPT TNC^Null^ tumors lacked TNC, despite the presence of vimentin+ stromal cells at the tumor edge (Fig. 3h and Supplementary Fig. 3e). Since LUADs are highly heterogeneous, we applied GLASS-AI^55^ to assess the proportion of the tumor presenting as the more aggressive grades 3 and 4. The most aggressive grade 4 designation was decreased in early tumors that lacked TNC (Fig. 3f, g). Staining for Proliferating Cell Nuclear Antigen (PCNA) showed cell proliferation at the tumor edge was decreased in the TNC^Null^ tumors (Fig. 3h, i). Quantification of the tdTomato+ tumor cells in the tumor microenvironment region outside of the main tumor mass confirmed that TNC^Null^ tumors harbor fewer invasive cells than TNC^WT^ tumors (Fig. 3j, k). When we let tumors develop in KPT and KPT TNC^Null^ mice for 15 weeks, we found that overall tumor burden was significantly decreased in the TNC^Null^ tumors compared to tumors with TNC (Fig. 3l, m and Supplementary Fig. 3f). Thus, TNC expression at the edge of transitioning LUAD drives tumor cell proliferation and aggressiveness that later results in higher tumor burden.

We investigated how TNC expression in early LUAD induces tumor progression. Since TNC stiffens glioma tissues^34^ and its expression during fibrosis is associated with lung stiffening^56^, we tested if TNC was associated with tissue rigidity in LUAD. We used atomic force microscopy (AFM) to measure the elastic modulus of the LUADs in lung tissue slices of KPT tumors after 15 weeks of tumor growth (Supplementary Fig. 4a). However, we did not detect tissue stiffening in early LUAD (Supplementary Fig. 4b). Furthermore, we did not observe gross changes in ECM structure in the tumor microenvironment (Supplementary Fig. 4c). We detected a minor increase in collagen fiber thickness, but no change in pore size in 10 week KPT LUAD compared to WT lung tissue, although both metrics were significantly increased in late stage, high grade tumors (Supplementary Fig. 4d, e). These data suggest that TNC induces early LUAD progression by a mechanism other than by controlling tumor microenvironment stiffness or structure.

In addition to affecting the structure and stiffness in the tumor microenvironment, TNC also directly binds and activates integrins^57^, which signal to FAK, ERK, and other growth factor-activated and mechano-signaling pathways^58^. We therefore tested if TNC activates integrin signaling in early LUAD. GSEA analysis found increased expression of two different focal adhesion gene signatures in LUADs with the highest *TNC* expression (Fig. 4a). Focal adhesions are integrin-containing structures that bind the ECM and signal via focal adhesion kinase (FAK) to promote cell proliferation, survival, and migration^59, 60^. TNC has been shown to directly bind integrins αvβ1, αvβ3, αvβ6, α8β1, and α9β1^61, 62^. Of these integrins, *ITGAV* and *ITGB1* are the most highly expressed integrin subunits in LUAD^63^. We used available blocking antibodies against integrin αv, β1, and αvβ6 and chemical inhibitors against αvβ1 and αvβ3 to test the role of these integrins in TNC signaling to LUAD tumor cells. Blocking αv and β1 and inhibiting αvβ1 dramatically reduced H1299 LUAD cell proliferation and migration on TNC (Fig. 4b, c, and Supplementary Fig. 4f). Disrupting αvβ3 or αvβ6 signaling also caused measureable, but minor reductions in migration velocity on TNC (Fig. 4c and Supplementary Fig. 4f). Since inhibiting αvβ1 caused the largest reduction in migration among the αv integrin dimers, we tested if TNC signals to tumor cell integrin β1 in the native lung tumor environment. We used the HUTS-4 antibody to detect the activated form of integrin β1 in precision cut lung slices (PCLSs) of lung tumors from KPT mice 10 weeks after tumor induction. Immunofluorescence staining and quantification of HUTS-4 intensity in tdTomato+ tumor cells and local TNC intensity showed that *in vivo* activation of tumor cell integrin β1 directly correlated with the cell’s local TNC level (Fig. 4d, e), consistent with TNC activating integrin β1 in early LUAD tumor cells.

**Figure 4.**
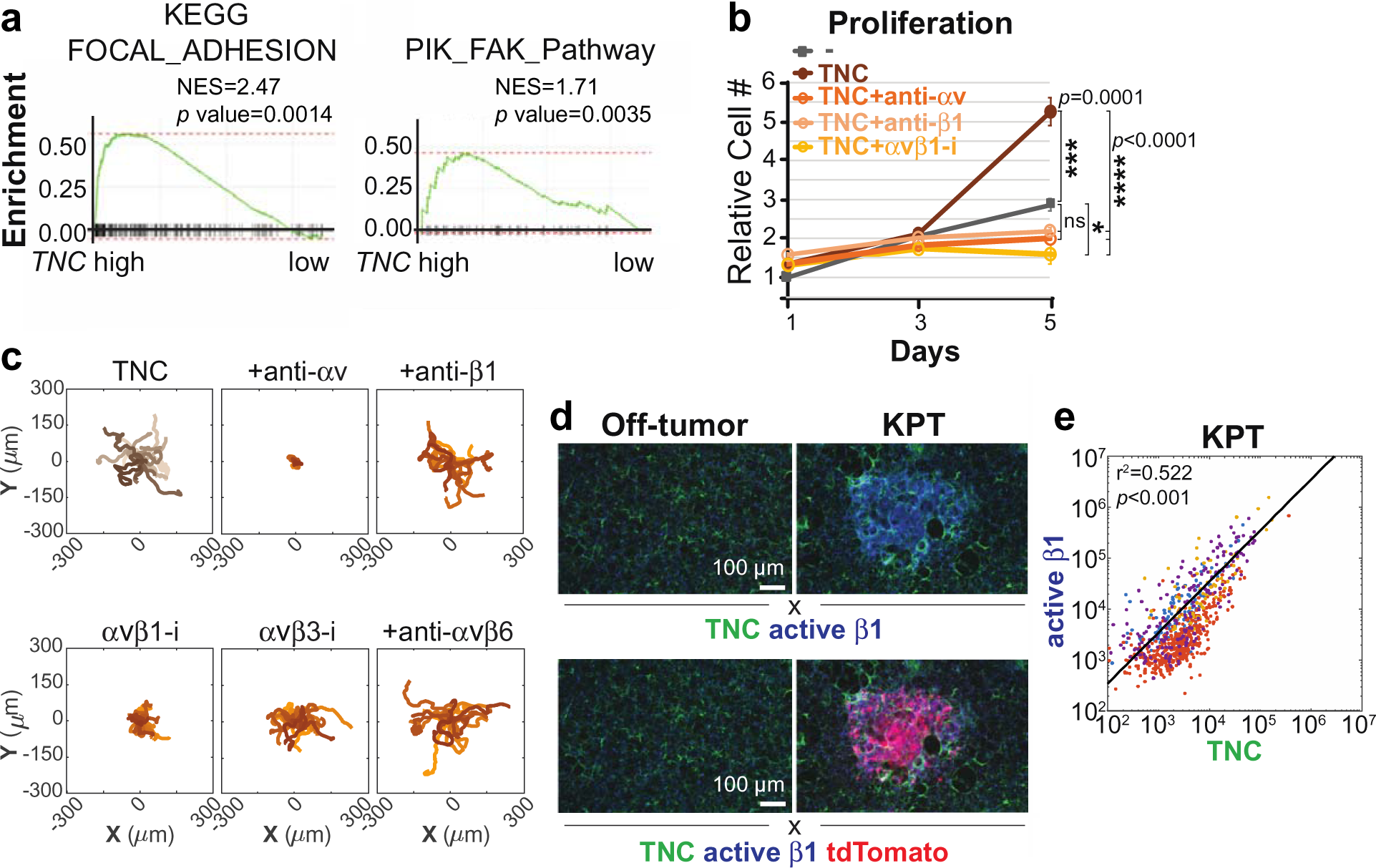
TNC signals to LUAD tumor cells through integrin αvβ1. **a** GSEA plots comparing expression of focal adhesion signatures in TCGA with the highest and lowest quartile *TNC* expression. **b, c** H1299 cell proliferation and migration with integrin inhibitor treatment. Proliferation error bars are SEM for *n*=3 independent experiments. One-way ANOVA with Tukey’s test. Migration tracks are 20 representative cells from *n*=3 experiments. **d** Representative 3D confocal scans of immunofluorescence for activated integrin β1 and TNC in PCLSs of KPT transitioning LUAD, 10 wks, *n*=2 females and 2 males. **e** Sum of integrin β1 pixels within and TNC pixels near the tumor cells in images in (d). Colors represent the 4 different animals. R^2^ and *p* value for linear regression model.

We next tested if TNC signals to FAK during early tumor growth. H1299 cells plated on TNC had increased phospho (p-) FAK, relative to cells plated on control albumin, and the p-FAK was reduced by disrupting integrin αv and αvβ1 signaling (Fig. 5a, b). Treating live PCLSs from KPT mice with the αv blocking antibody reduced the levels of activated integrin β1 and p-FAK within the tumor cells, indicating that integrin αvβ1 signals to FAK within the native lung tumor setting (Fig. 5c, d). Inhibiting FAK reduced the baseline and TNC-induced proliferation of H1299 (Fig. 5e). As expected for a critical component of focal adhesion regulation^64^, inhibiting FAK blocked the baseline migration of H1299 cells on BSA and also reduced the TNC-induced migration (Fig. 5f and Supplementary Fig. 5). Immunohistochemistry for p-FAK in KPT TNC^WT^ and TNC^Null^ lungs showed p-FAK at the invasive edge of LUADs after 10 weeks of tumor growth, which was reduced in the TNC^Null^ tumors (Fig. 5g, h). The cells with high p-FAK were more pleomorphic, compared to cells with low p-FAK, exhibiting irregular cell and nuclear shapes and sizes that suggest an invasive transition^65^ (Fig. 5g). Multiplex immunofluorescence for p-FAK and TNC in clinical samples of stage I and II LUAD showed p-FAK levels in LUAD tumor cells were higher in the TNC-positive invasive regions versus TNC-negative regions (Fig. 5i, j). Thus, TNC expression in the tumor microenvironment is associated with increased FAK activation in the tumor cells and an aggressive tumor cell state. We then tested if early FAK activation signals for tumor progression by treating mice harboring 10 week KPT tumors with low dose FAK inhibitor (VS-4718) for 5 weeks. FAK inhibitor treatment significantly reduced the tumor burden present at 15 weeks, compared to control mice (Fig. 5k, l), indicating thta FAK signaling in early transitioning LUAD contributes to the tumor growth and progression.

**Figure 5.**
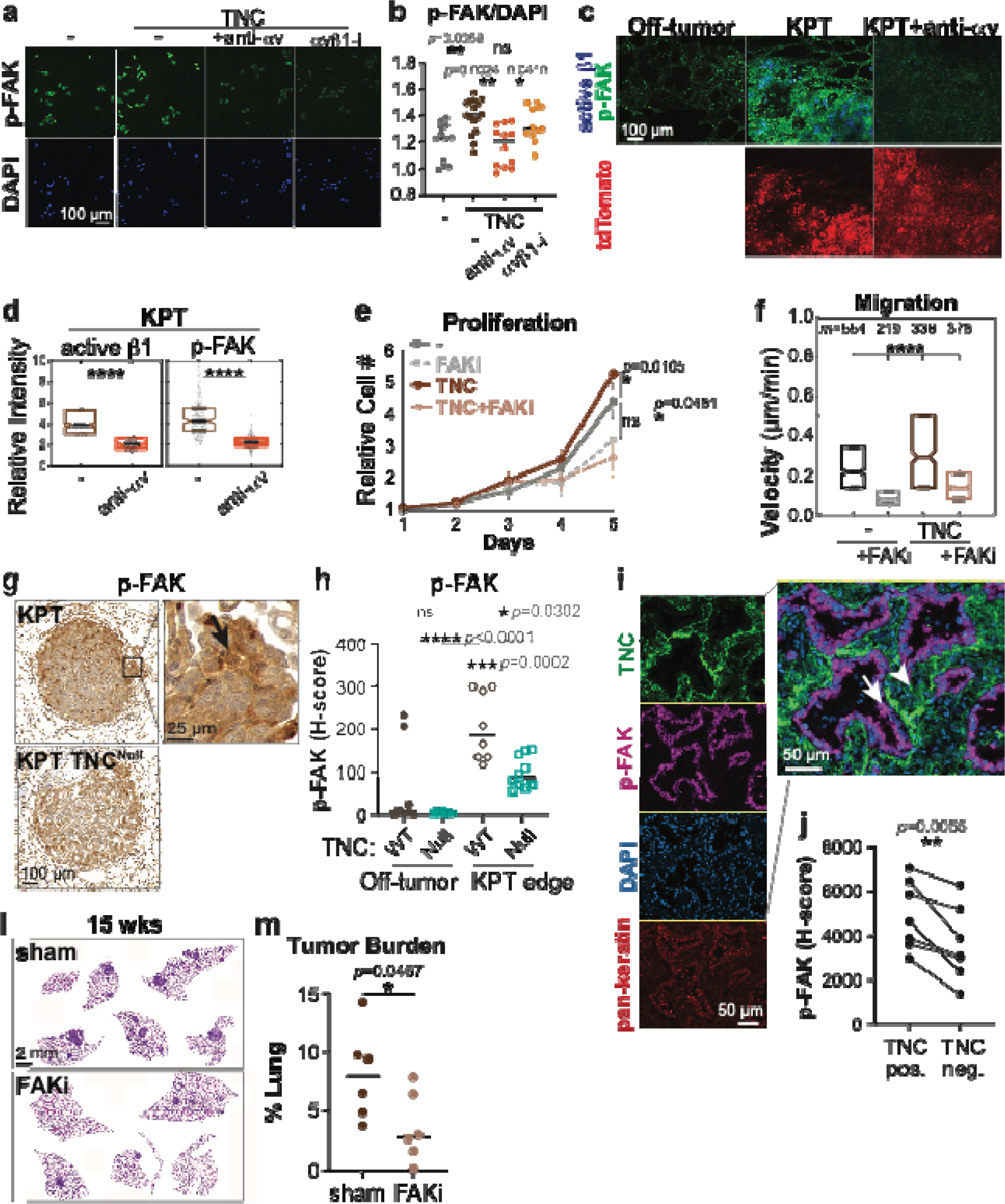
TNC signals to FAK in early LUAD. **a, b** Representative images and quantification of immunofluorescence for p-FAK in H1299 cells plated on BSA or TNC and treated with integrin inhibitors. Per field of view, total p-FAK intensity normalized to DAPI. *n=*3 experimental replicates, 3 fields of view per replicate. **c, d** Representative 3D confocal scans and quantification of active integrin β1 and p-FAK in immunofluorescence in PCLS from KPT 10 wk transitioning LUADs treated with and without integrin αv inhibitor. Plots show relative mean intensities of active integrin β1 and p-FAK in tdTomato+ tumor cells, with the median intensity labeled in black and 25^th^-75^th^ percentiles outlined in boxes. *n*=1 female and 1 male mouse. Mann-Whitney U test. **e, f** H1299 cell proliferation and migration with DMSO or FAKi PF-573228 treatment. Proliferation error bars are SEM for *n*=4 independent experiments. One-way ANOVA with Tukey’s test. Migration is *m* cells from *n*=3 independent experiments. K-S test. **g, h** Representative immunohistochemistry and quantification of p-FAK in KPT LUAD, *n=*2 female and 2 male mice of each genotype. Arrow marks tumor cells with p-FAK and pleomorphic shape. Arrowhead marks tumor cells without p-FAK and uniform shape **i, j** Representative multiplex immunofluorescence and quantification for p-FAK in TNC-positive and -negative regions in human LUAD. Six stage I and II LUADs selected as having moderate or high TNC from Supplementary Fig. 1j and invasive areas identified by a board-certified pathologist. Arrow marks tumor cells with high p-FAK, near TNC. Arrowhead marks tumor cells with low p-FAK, not adjacent to TNC. Two-tailed paired T-test. **k** Representative H&E image of KPT lung lesions, 15 weeks after Cre inoculation and treated with sham or FAKi VS-4718 for 5 weeks, *n*=3 female, 3 male mice for each treatment. **l** Quantification of tumor burden for (k). Unpaired T-test.

### Lasting TNC expression in the lung can contribute to tumor progression

Given that lung injury induces acute TNC expression^22, 25, 29, 30^ and TNC promotes the malignant progression of LUAD cells (Fig. 3 and Supplementary Fig. 3), we hypothesized that TNC could promote adaptive oncogenesis. If this were true, people with elevated pulmonary *TNC* expression would have greater risk of developing early LUAD and greater risk of locally recurrent LUAD following standard lobectomy for stage I/II cancers. We first tested if lung injury or advanced age resulted in sustained TNC expression. We exposed young mice to repeated low-dose bleomycin to model lung injury due to environmental exposures. 3 weeks after the initial dose and 1.5 weeks after the last treatment, the mouse lungs exhibited significant expression of TNC (Fig. 6a, b). Male mice carried 2 weeks further past the last treatment still showed pockets of lasting TNC in their lungs (Fig. 6a). Interestingly, the sustained TNC expression after injury was unique to the male mice (Fig. 6c). We also analyzed the lungs of transgenic mice that harbor a tamoxifen-inducible Surfactant Protein C^I73T^ (SPC^I73T^) mutation, which induces lung injury^66^. We compared TNC levels in the lungs of uninduced mice at different ages. Lungs from old mice (2 years old, ∼80 human years) exhibited enlarged alveoli due to age-induced elastin degradation^23^ and irregular TNC expression that was not present in lungs from young adult mice (2-3 months of age) (Fig. 6d). However, the frequency of TNC-positive alveoli was too low to affect overall TNC levels in the aged lung tissue (Fig. 6e). We then induced lung injury in young adult mice with tamoxifen treatment, which caused uniform TNC expression at 2 weeks. By 6 weeks, the initial injury response resolved, but residual alveoli with thickened walls and TNC expression remained (Fig. 6d, e). This suggested that during one’s lifetime, exogenous stressors could result in a cumulative burden of TNC expression that leads to alveoli with greater propensity of developing invasive LUAD.

**Figure 6.**
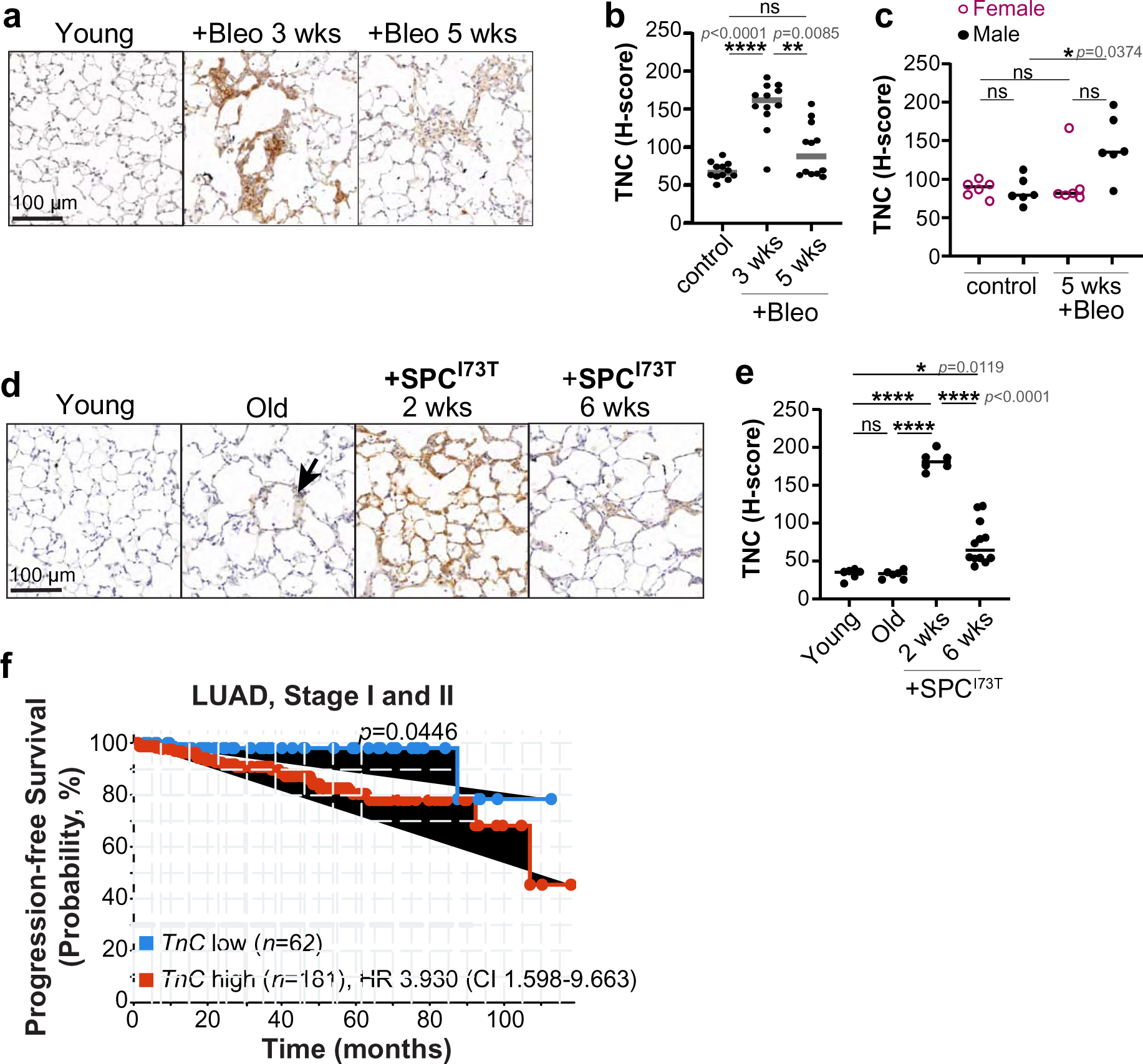
TNC is expressed in response to environmental stressors and predicts LUAD progression and recurrence. **a-c** Representative immunohistochemistry and quantification of TNC in C57Bl/6J mice. Young controls (60-90 days) and young treated with bleomycin (4 doses of 0.25 mg/kg) and assayed 3 or 5 weeks after the initial dose. *n*=2 female and 2 male mice per condition. Kruskal-Wallis test. **d, e** Representative immunohistochemistry and quantification of TNC in *SPC^I73T^*mice before and after induction with tamoxifen. *n*=2 female and 2 male mice for control young, *n*=3 female and 3 male for old, *n*=4 female and 4 male with 2 weeks of *SPC^I73T^* expression, and *n*=2 female and 2 male with 5 weeks of *SPC^I73T^* expression. One-way ANOVA with Tukey’s post-hoc test. **f** Recurrence-free survival in 243 stage I and II LUAD patients without distant metastasis from ORIEN network. Patients are split into lower 25% (≤12.85 normalized Log2 fold change) and upper 75% (>12.85) of *TNC* expression. Log-rank test. HR is Hazard Ratio with 95% CI, Confidence Interval.

We tested if *TNC* expression is associated with the recurrence of early LUAD using clinical samples of stage I/II. We obtained tumor RNA-sequencing data and clinical outcomes for >500 stage 0, I, and II LUAD patients in the ORIEN network of NCI-designated comprehensive cancer centers (https://www.orientcc.org). We limited our analysis to samples with ≥ 10% of non-tumor cell RNA, to ensure that we captured the tumor microenvironment. Stage I/II LUAD patients are treated with a partial or complete lobectomy of the lung lobe bearing the tumor and in some cases, additionally with adjuvant chemo-, immune-, or targeted therapy. We followed patients’ outcome beginning 45 days after their surgery and identified patients with no progression or recurrence and a group of patients with local cancer recurrence or metastases within the lung. Patients with progression due to distant metastasis were excluded. In this population, patients whose tumor samples harbored higher *TNC* expression experienced a shorter time to LUAD progression, compared to those with low *TNC* expression (Fig. 6f). High *TNC* conferred a hazard ratio of 3.93, meaning that these patients have nearly 4 times the probability of recurrence, compared to patients with low *TNC*. Since stage is also prognostic, we repeated the comparison with patient samples that were unambiguously stage I or stage II. While the sample size was not powered to detect a significant different in outcome, both tests showed the same trend of shorter time to LUAD progression in samples with less *TNC* expression (Supplementary Fig. 6). This association suggests that *TNC* expression promotes the transition of pre-existing *in situ* lesions into invasive LUAD, although progression due to the outgrowth of undetected tumor cells from unresected lymph nodes cannot be excluded.

## Discussion

The role of ECM changes in the acquisition and progression of early LUAD is poorly understood. Early LUAD lacks the hallmarks of late stage metastatic LUAD, such as a complex genetic landscape and desmoplastic stroma. Yet, a high rate of recurrence persists and drives mortality^3, 67^. We show here that production of TNC by fibroblasts can push early tumors towards progression and recurrence. Our finding that exogenous stressors can cause TNC accumulation further suggests that TNC could function in adaptive oncogenesis to promote the development of lung cancer.

We identified a TNC→integrin αvβ1→FAK pathway activated at the tumor edge that promotes LUAD cell proliferation and migration. We observed increased TNC at the edge of early tumors in transgenic models with the *Kras^G12D^* driver mutation alone and with *Trp53* or *Lkb1* loss, and also in orthotopic transplants with *Nkx2-1* loss. While TNC induces glioma tissue stiffening, which correlates with tumor aggressiveness^34^, its expression at the edge of early lung tumors was not associated with tissue stiffening. Instead, TNC expression at the edge of both mouse and human tumors was associated with signaling to nearby tumor cells through β1 integrin and FAK activation. Tumors in mice lacking TNC exhibited reduced p-FAK, reduced tumor cell proliferation and invasion, and reduced area of high-grade pathology, when compared to mice with TNC. These findings are consistent with recent studies showing that FAK drives tumor cell aggressiveness. *Cdkn2A* and *Lkb1*-mutant LUAD mouse models produce high-grade tumors with high FAK activation, compared to *Trp53*-mutant tumors^13, 68^. The high FAK activity is required for the tumors’ aggressive phenotype and especially present in foci of invading cancer cells^13, 68^. While FAK activation is generally low across human LUADs, activated FAK increases in and promotes residual disease after targeted therapy against RAS-pathway oncogenes^69, 70^. The TNC-associated FAK activation develops at a time when a high-plasticity cell state emerges, which is also associated with high tumor burden, drug resistance, and poor patient prognosis^10^. We show that inhibition of this FAK activity in established tumors reduces early tumor burden. Thus, we propose that in early tumors with *RAS* oncogenes *or RAS* and *TRP53*-mutations, TNC signaling creates pockets of tumor cells with high FAK activity that drive tumor progression. For patients with high TNC expression, treatment with FAK inhibitors could help prevent tumor progression.

TNC is induced by fibroblasts in early tumors and additionally in tumor cells in advanced, high-grade LUADs. We show that activated, αSMA-positive fibroblasts in the tumor microenvironment are the main source of TNC in the earliest LUADs modeled in the KPT mice. Such activated fibroblasts are on a trajectory to transition into myofibroblasts, which correlates with poor LUAD survival^71^. Interestingly, as the tumors progress into early cancers, the tumor cells themselves produce TNC. In our and others’ mouse models of metastatic, high grade LUAD with desmoplastic stroma, TNC expression occurs throughout the tumor and tumor microenvironment, driven by tumor cell loss of NKX2-1^30^. In human stage I/II LUAD, which have not spread to distant lymph nodes or other parts of the body and retain some NKX2-1 expression, we show that both fibroblasts and tumor cells express TNC. LUAD tumors cells likely begin to acquire *TNC* expression as they develop plasticity, since the high plasticity cell state is characterized by low *NKX2-1* expression^10^. This suggests that the earliest tumors create a wound environment that activates local fibroblast at the tumor-stroma interface. These TNC-producing activated fibroblasts could be a new therapeutic opportunity for preventing early lesions’ transition to aggressive cancer or recurring after surgery.

The long delay between the appearance of oncogenic mutations and LUAD development^5, 8, 9^ and the high recurrence rate of stage I/II cancers suggest that exogenous stressors contribute to cell transformation and cancer progression. Lung injury induces a repair process involving transient fibroblast activation and ECM production^22, 25, 42^. Age also causes fibrotic changes in the lung, including increased fibroblast number and altered ECM^72^. We show that these stressors induce pockets of persistent TNC expression in the lung. We also observed greater TNC expression in response to lung injury, as well as in adenomas and LUADs, in males versus females, suggesting that male sex could further contribe to TNC expression. Our analysis of clinical samples of stage I/II LUAD showed that *TNC* expression is associated with recurrence in patients. Thus, we propose a model in which benign or early cancerous lesions that result from tumor-initiating mutations like oncogenic *KRAS^G12D^* are activated for progression by exogenous stressors that cause pockets of sustained TNC, which signal to the neighboring pre-cancer or early cancer cells for FAK activation and aggressiveness. Pollution and fibrosis have also been shown to increase primary lung tumor growth and metastatic seeding of LUAD cells in the lung by affecting immune cell recruitment^24,73^. Thus, these stressors likely act through multiple mechanisms, including fibroblast activation and direct ECM signaling to tumor cells. Fibroblast activation and high TNC expression could identify early LUAD patients that would benefit from adjuvant therapy after surgery to prevent recurrence.

## Abbreviations

ECM: extracellular matrix
FAK: focal adhesion kinase
GEMM: genetically-engineered mouse model
LUAD: lung adenocarcinoma
TNC: Tenascin-C

## Supplementary Figures and Tables

**Supplementary Figure 1.**
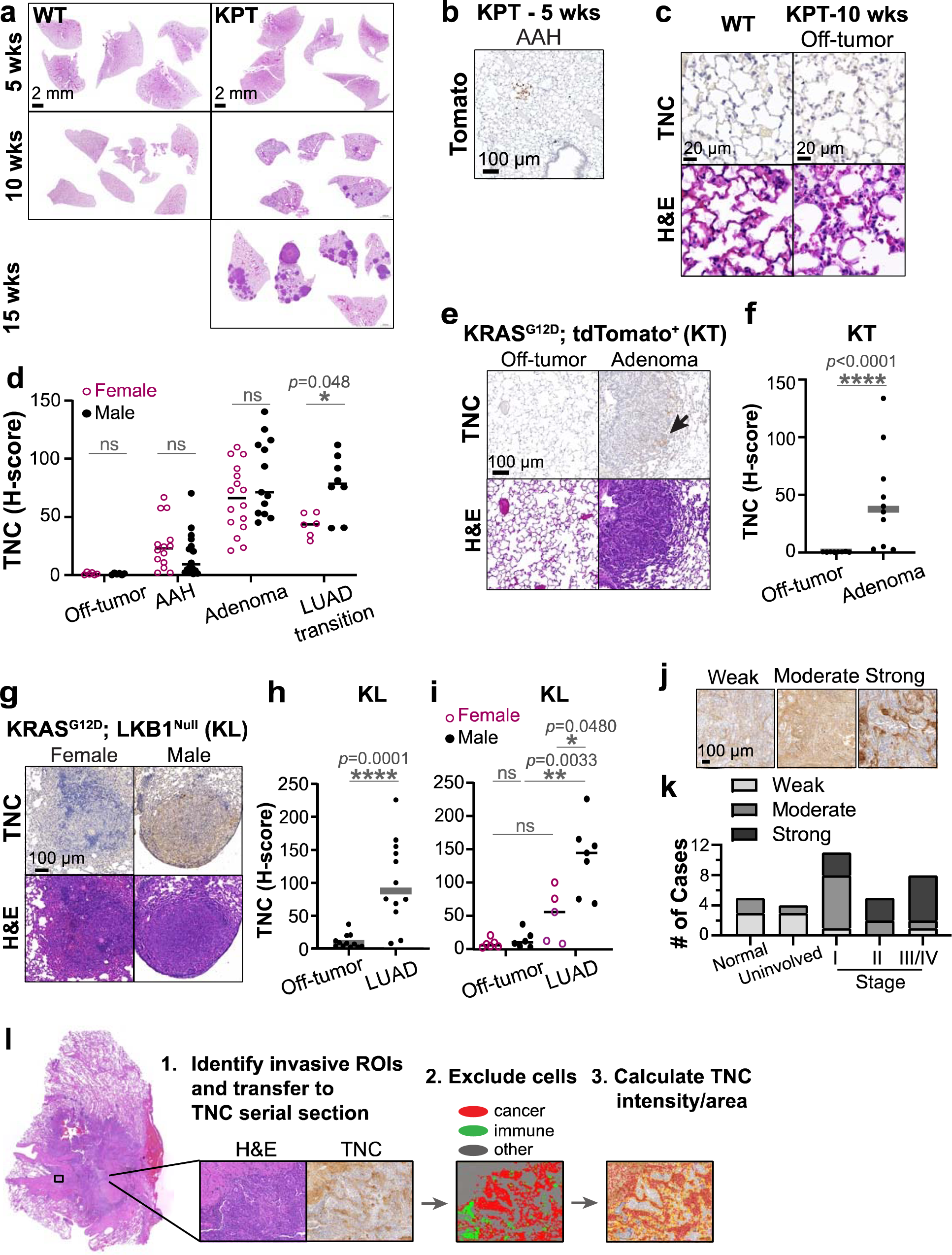
TNC is expressed at the tumor-stroma interface in early LUAD. **a** Representative full scans of H&E-stained lung lobes after Cre inoculation. KRAS WT lungs from *Kras^+/+^; Trp53^f/f^; tdTomato* mice, *n*=2 mice at 5 wks, *n*=3 at 10 wks, and *n*=6 at 15 wks. KPT tumors from *KRas^LSL-G12D/+^; Trp5^F/F^; Rosa^LSL-tdTomato^* mice, representative of *n*=2 mice at 5 weeks and *n*=10 at 10 weeks and 15 weeks. **b** Representative immunohistochemistry for tdTomato in KPT lungs, 5 weeks, *n*=2 mice. **c** Representative immunohistochemistry for TNC and H&E stain for lung structure in WT lungs and off-tumor region of KPT lungs, *n*=3 mice for each genotype. **d** Quantification of TNC in lesions from KPT lungs in Fig. 1b by sex. Two-way ANOVA with Sidak’s multiple comparison test. Dots are individual H-scores for each Region of Interest (ROI) and lines are median. **e-h** Representative immunohistochemistry for and quantification of TNC in adenomas from KT mice (*n*=2 females, 2 males) and low grade tumors from KL mice (*n*=3 females, 3 males). Arrow marks low TNC staining. Mann Whitney test for KT and one-way ANOVA with Tukey’s posthoc test for KL. **i** Quantification of TNC by sex. Two-way ANOVA with Sidak’s multiple comparison test. **j, k** Representative images and tabulation of samples with low, moderate, and high TNC in human LUAD and normal lung tissue in Fig. 1j. **l** Machine learning workflow used in Fig.1k to automatically identify and quantify stromal TNC in off-tumor and invasive LUAD ROIs. ROIs from the H&E stain were transferred to immunohistochemistry stain for TNC, tumor and immune regions were excluded, and TNC/area within each ROI was calculated. In step 3, Blue indicates negative for TNC, Yellow low TNC, Orange medium TNC, Pink high TNC.

**Supplementary Figure 2.**
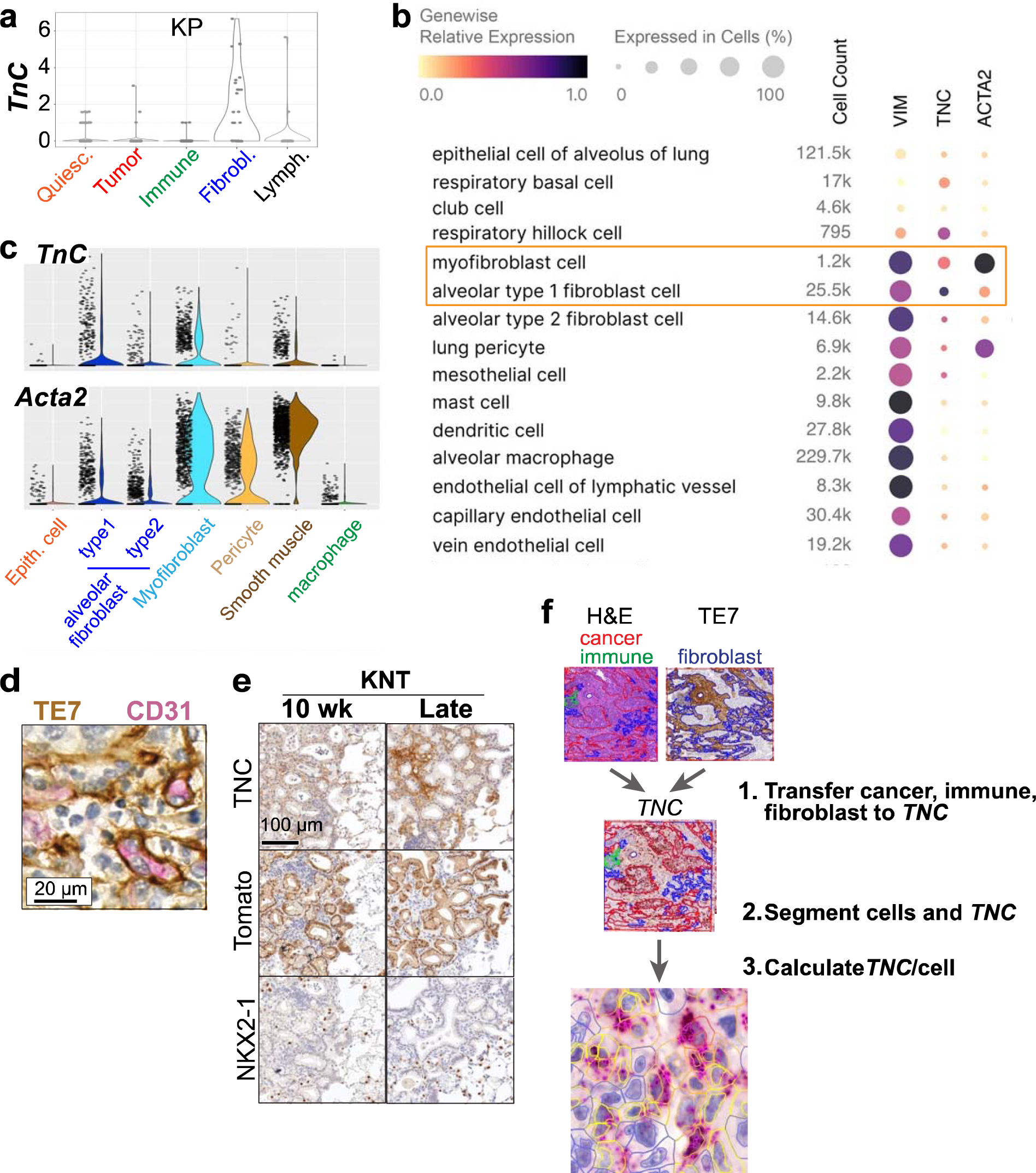
TNC is expressed by fibroblasts at the tumor-stroma interface in mouse tumors transitioning to LUAD and stage I and II human LUAD. **a** Plot of log2 normalized counts of *TnC* in each cell types identified in scRNAseq of KP tumors^41^. **b, c** Analysis of *TNC* and fibroblast marker expression in HLCA^42^. **d** Sequential immunohistochemistry co-detection of TE7+ cells and CD31+ endothelial cells in human LUAD, to evaluate fibroblast specificity for TE7, representative of *n*=3 cases. **e** Immunohistochemistry for TNC and tumor markers in early and late tumors from *KRas^LSL-G12D/+^;Nkx2-1^F/F^; Rosa^LSL-tdTomato^*(KNT) mice. Representative of tumors from *n*=3, 2, and 2 male mice, respectively. **f** Workflow of automated quantification of *TNC* puncta in cancer, immune, and fibroblast cells. In step 3, Blue indicates negative for *TNC*, Yellow 1+ *TNC*/cell, Orange 4+ *TNC*/cell, Pink 10+ *TNC*/cell.

**Supplementary Figure 3.**
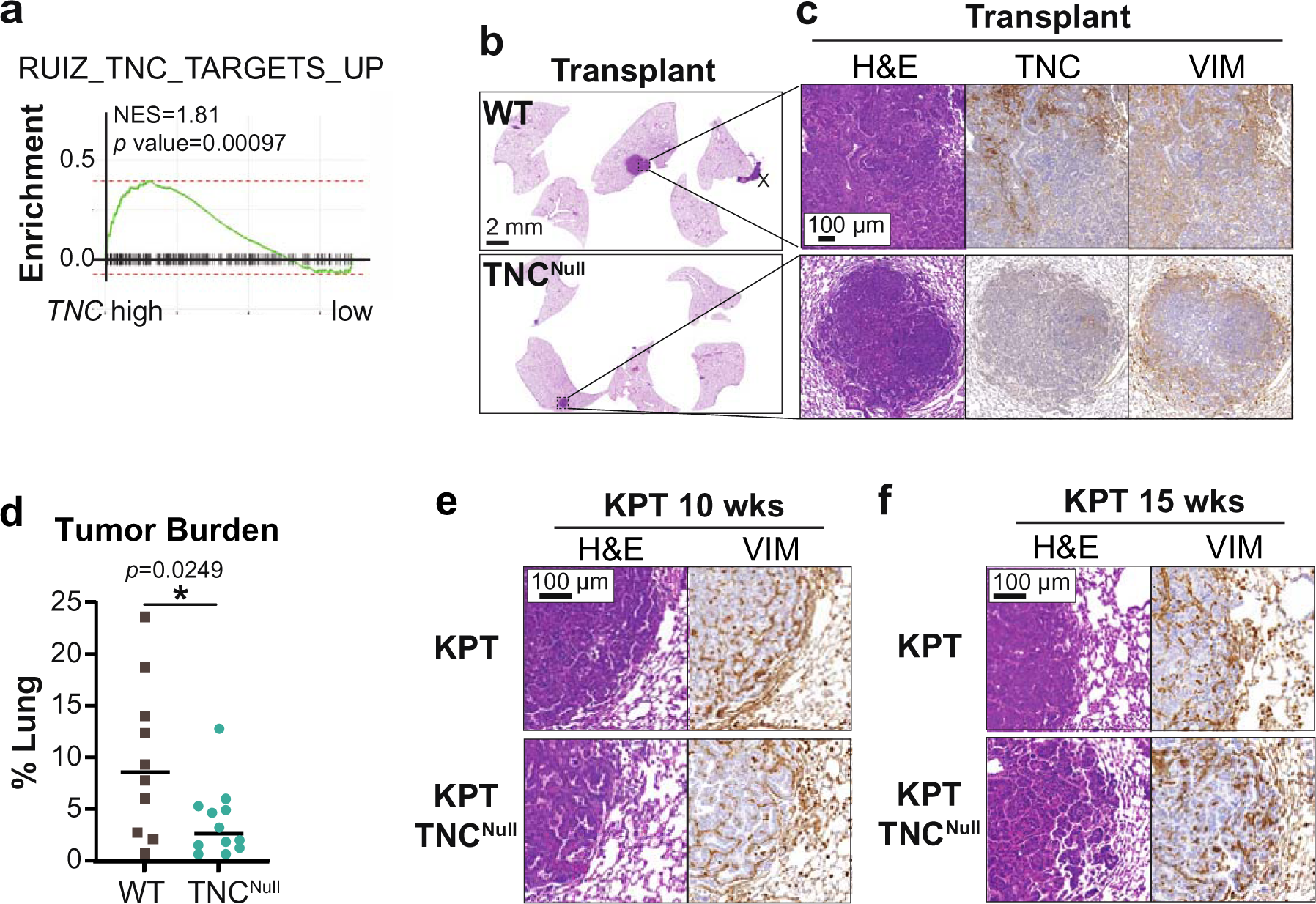
TNC promotes early LUAD progression. **a** Gene set enrichment analysis (GSEA) plot comparing expression of TNC-response genes between LUADs in TCGA with the highest (top quartile) and lowest (bottom quartile) *TNC* expression. NES is normalized enrichment score. **b** Representative low magnification lung lobe overview with H&E staining of lung tumors from orthotopic transplant of 3658 cells. Tumor cells that grew on the outside lung surface were excluded from the analysis, marked with X. **c** Representative high-magnification view of H&E stain in (b) and immunohistochemistry for TNC and Vimentin in serial sections, *n=*5 female and 5 male TNC^WT^, *n=*6 female and 6 male TNC^Null^ mice. **d** Tumor burden per mouse, with tumor >25,000 μm^2^. Mann-Whitney U test. **e, f** Representative H&E stain and immunohistochemistry for Vimentin on serial sections of KPT lungs 10 weeks and 15 weeks after tumor induction, from Fig. 3d, l.

**Supplementary Figure 4.**
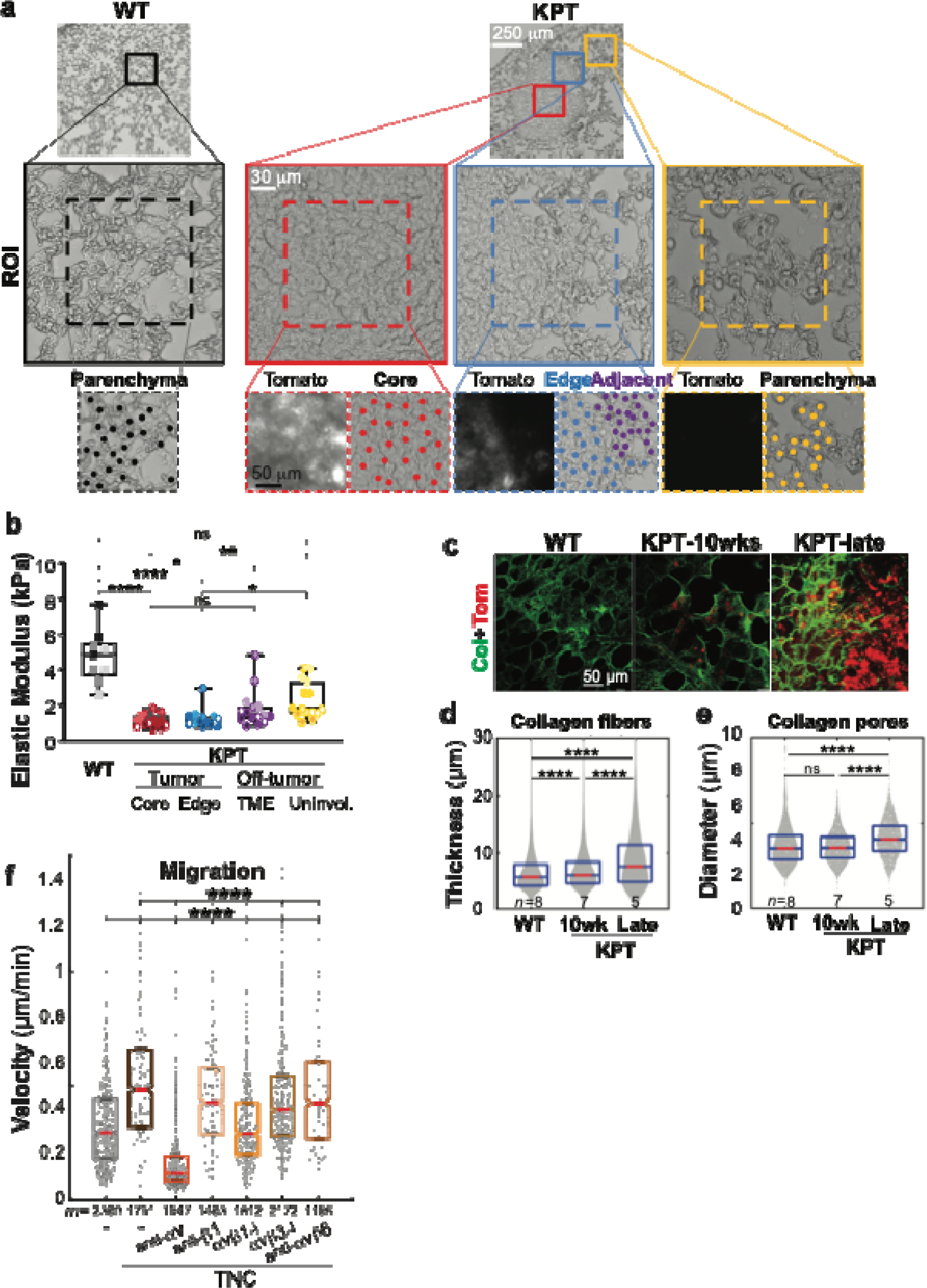
TNC in LUAD is not associated with ECM stiffening or bundling in early LUAD. **a, b** AFM workflow and quantification of LUAD tumor tissue, shaded points indicate 3 measurements for each region, with samples from *n*=5 WT and *n*=5 KPT mice 15 weeks after inoculation with Adeno-Cre. TME is tumor microenvironment. **c** 3D confocal scans of PCLS from WT lungs and KPT LUAD, with collagen (Col) labelling with CNA35-GFP and tdTomato (Tom) by immunofluorescence. 3 ROIs imaged for each mouse, with *n*=3 WT mice, 3 KPT mice at 15 weeks and 2 KPT mice at 26 weeks, with at least 1 of each sex. **d, e** Quantification of collagen fiber thickness and pore diameters in (c). **f** H1299 cell migration velocity with treatment with integrin inhibitors. *m* cells from *n*=3 independent experiments in (4c).

**Supplementary Figure 5.**
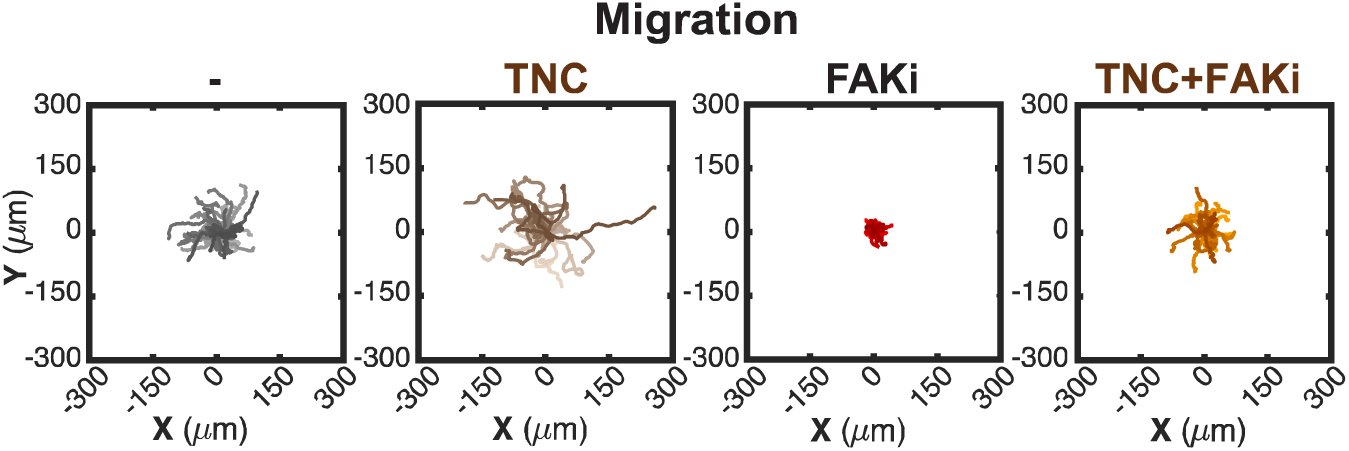
FAK is required for LUAD cell migration. H1299 cell migration on plates coated with BSA or TNC and treated with FAK inhibitor. Rose plots show tracks of 40 representative cells in Fig. 5f.

**Supplementary Figure 6.**
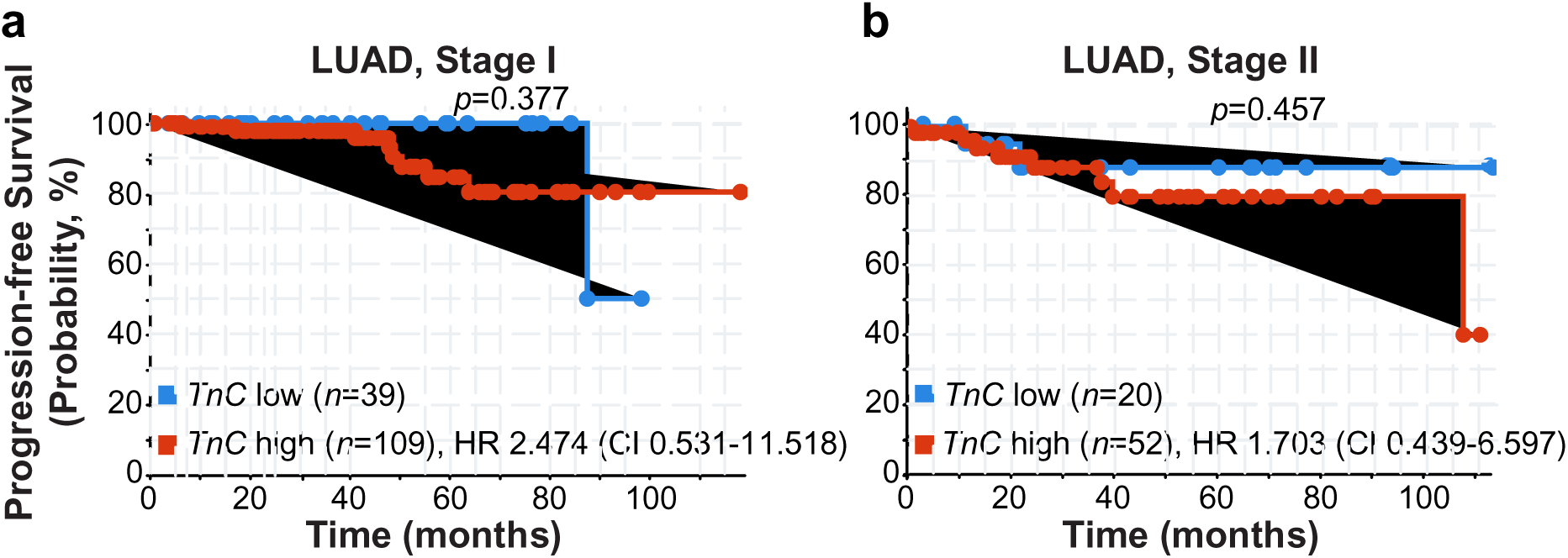
Patients with low TNC trend to have a lower risk of recurrence. **a** Recurrence-free survival of stage I and **b** stage II LUAD patients in Fig. 6f, separated by stage determined by the ORIEN network Pathology group. Patients are split into those with ≤12.85 normalized Log2 fold change and >12.85 of *TNC* expression. Log-rank test. HR is Hazard Ratio with 95% CI, Confidence Interval.

**Supplementary Table 1.**
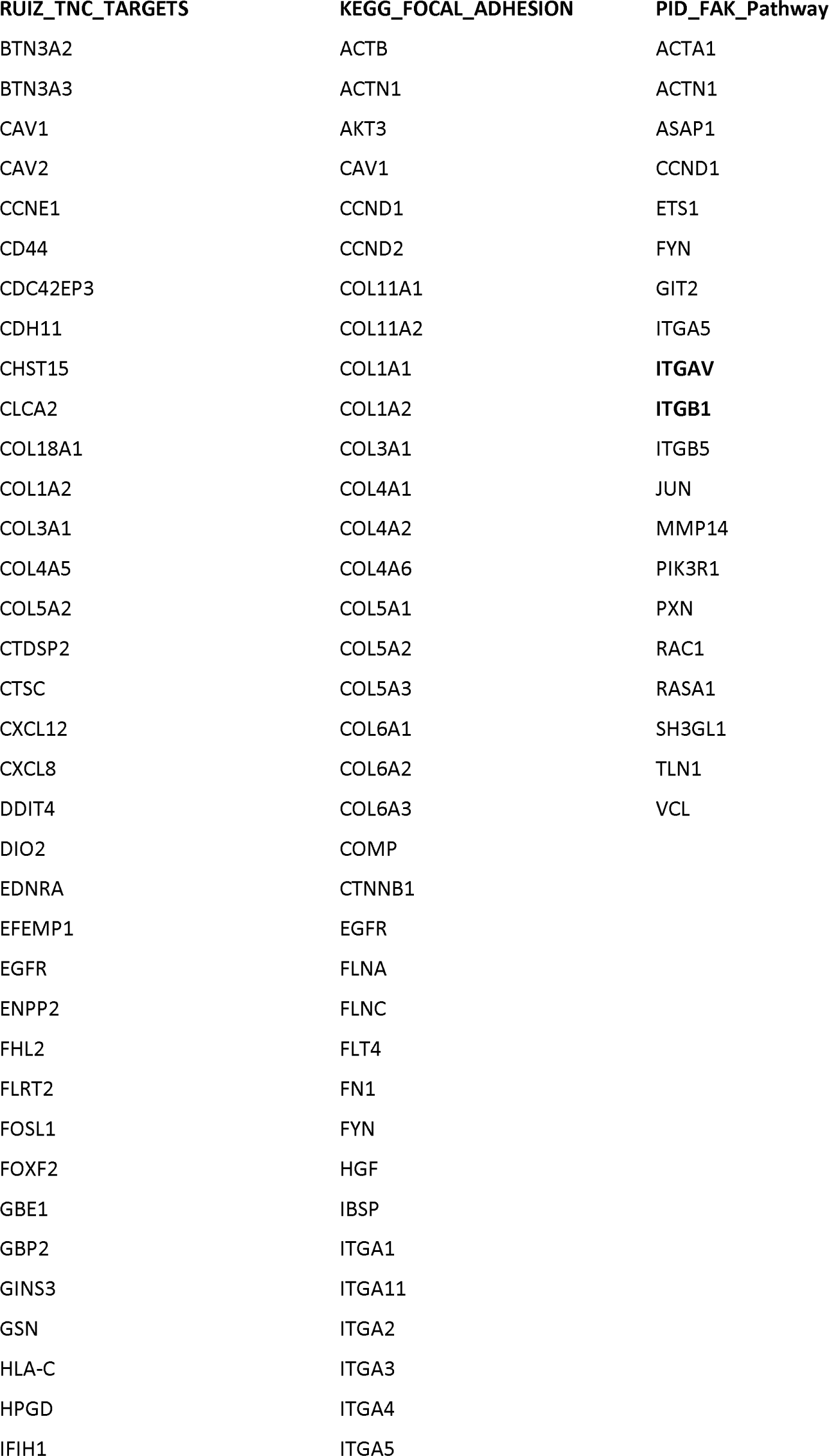
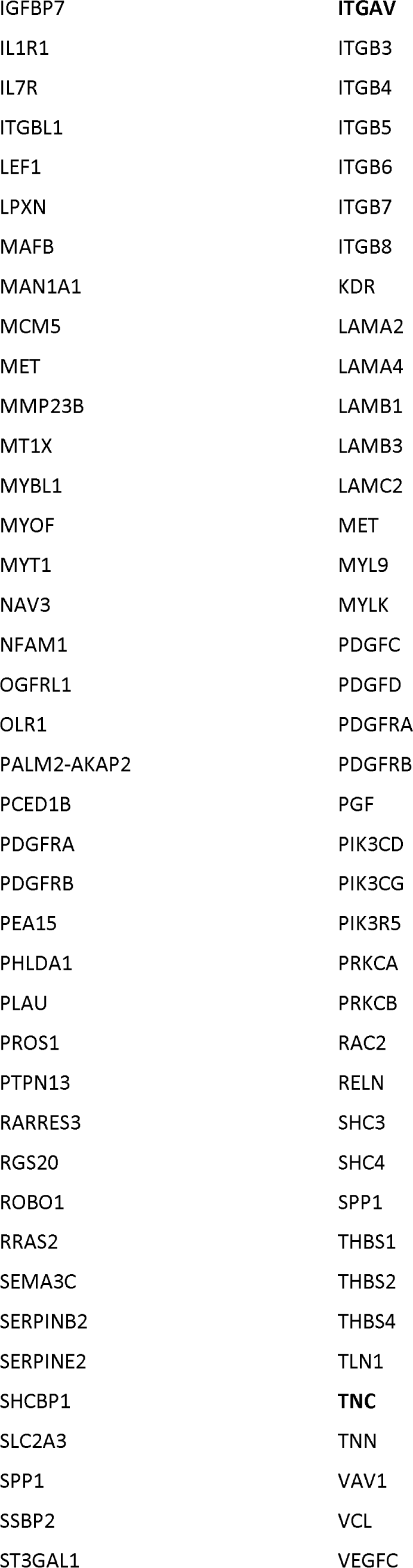
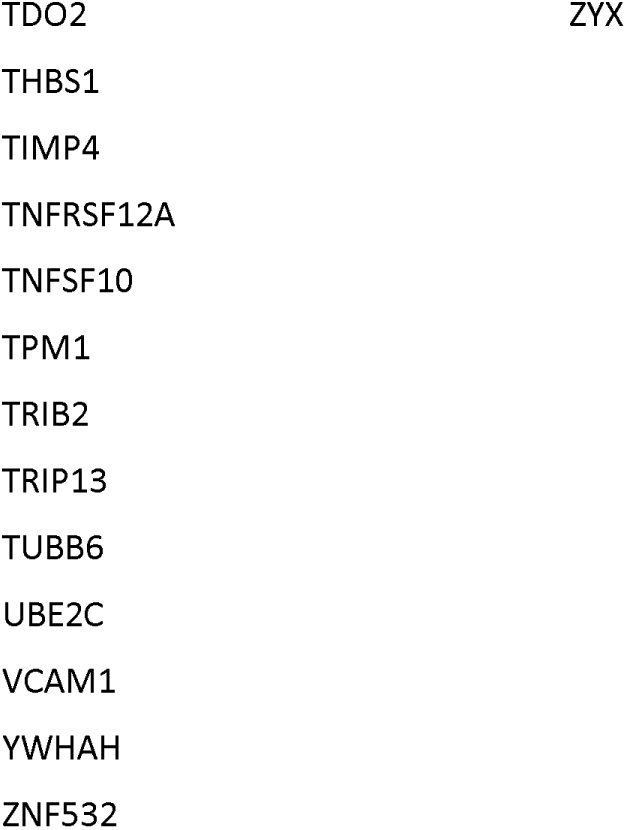
Core genes (leading edge genes) expressed by TNC-high samples that drive their enchrichment of TNC Targets and Focal Adhesion gene signatures.

**Supplementary Table 2.** Gene ontology (GO) terms for core TNC target genes.

| Fold Enrichment | Pathways |
| --- | --- |
| 7 | cellular response to growth factor stimulus |
| 6.8 | response to growth factor |
| 6.6 | response to wounding |
| 5.6 | regulation of cell migration |
| 5.2 | regulation of cell motility |
| 5 | regulation of locomotion |
| 5 | circulatory system development |
| 4.9 | regulation cellular component movement |
| 4.4 | cell migration |
| 4.3 | response to endogenous stimulus |
| 3.9 | cell adhesion |
| 3.9 | cell motility |
| 3.9 | localization of cell |
| 3.7 | locomotion |
| 3.3 | cell surface receptor signaling pathway |
| 3.2 | anatomical structure morphogenesis |
| 3.1 | cellular response to organic substance |
| 3.1 | response to organic substance |
| 2.4 | system development |
| 2.4 | response to chemical |

## Ethics Statement

Human data collected and analyzed for this research project was approved by the University of Utah Institutional Review Board (approval #00141909I/F and 89989). Prior to obtaining tissue for analysis, all samples were de-identified to comply with HIPAA regulations. The staining and analysis of slides were carried out in accordance with relevant guidelines and regulations.

## Acknowledgements

The research reported herein utilized the Huntsman Cancer Institute’s Preclinical Research Resource and Biorepository and Molecular Pathology (BMP) Shared Resources, as well as the Research Informatics, Research Histology, Quantitative Pathology, and Bioinformatics Cores and the University of Utah Cell Imaging Core. We acknowledge the University of Virginia, Charlottesville as a member of ORIEN and Verastem for their generous gift of VS-4718. The research was funded by grant #RSG CSM-130435 from the American Cancer Society, the Concern Foundation, and CA255790 from the National Cancer Institute of the National Institutes of Health. We acknowledge additional direct financial support for the research by the Huntsman Cancer Foundation and the Cell Response and Regulation Program at the Huntsman Cancer Institute and support by the National Cancer Institute under Award Number P30CA042014. The content is solely the responsibility of the authors and does not necessarily represent the official views of the NIH, the Department of Defense, or the U.S. government.

## Methods

### Kras-driven Mouse Models of LUAD

Mice were maintained under the University of Utah IACUC guidelines under protocols 18-08005, 21-10007, and 00-001500 and Emory University PROTO201700269. The *KRas^LSL-G12D/+^;Trp53^F/F^;Rosa26^LSL-tdTomato^* (*KPT*) and (*KNT*) strains were maintained on a mixed C57BL/6J 129SvJ background. To induce tumors, mice 55-105 days old were intratracheally infected with 1×10^8^ pfu/mouse SPC-Cre Adenovirus. Early tumors from mice terminated 5, 10, or 15 weeks after Cre inoculation and late tumors from mice terminated 20-26 wks. ∼50% of mice had tumors at the early time points. Off-tumor was ≥ 10 mm from tdTomato+ cells, adenoma <200,000 μm^2^ with at least one filled-in alveolus, transitioning LUAD at 10 weeks >200,000 μm^2^ with disrupted alveolar architecture with invasive cells outside of the main tumor mass. Representative IHC and IF images are from male mice unless otherwise noted.

Slides from KL tumors were from the *Rosa26^LSL-luciferase^; Kras^LSL-G12D/+^; Lkb1^F/F^ (KL_Luc_)* mouse intratracheally infected with Lentivirus-CMV-Cre-GFP-Puro and harvested within 12 weeks of tumor progression. Tumor grade was confirmed to be early-stage by a veterinary pathologist through H&E staining.

For FAK inhibitor treatment, KPT mice inoculated with Cre were microCT scanned at 10 weeks. Mice with notable lung lesions were enrolled in either a sham or VS-4718 treatment group by an investigator blinded to their tumor size. Mice were treated with saline or 50 mg/kg VS-4718 once daily by oral gavage, with a hiatus on the weekends, for 5 weeks.

### Lung injury and orthotopic tumor models

3658 LUAD cells were mixed with Matrigel and surgically transplanted into the lungs of *TnC* WT (C57BL/6) or *TnC* KO (C57BL/6) mice. 1×10^6^ 3658 cells were injected alone, or 9×10^5^ 3658 cells were co-injected with 2×10^5^ fibroblasts. Lungs were harvested 4 weeks after injection. In general, each *TnC* WT and *TnC* KO animal harbored one primary tumor.

Mice were 50-90 days old (young) and 560-690 days old (old). For lung injury, young mice were treated with 0.25 mg/kg of bleomycin 4 times, once every 4 days, by intratracheal intubation. *SPC^I73T^* mice were induced as previously described^66^.

### Human LUAD

Human data collection, de-identification, and analysis was approved by the University of Utah IRBs 00141909I/F and 89989. Tissue blocks from autopsies and LUAD lung resections were selected by the board-certified study pathologist (L.L.E) based on stage, size, histopathological feature, and tissue preservation. Autopsy cases were limited to patients with no history of cancer, lung infection, or chronic lung disease. Regions of invasive cancer were identified from H&E sections. “Uninvolved lung” was >4 cm from any tumor border and histologically benign. “Off-tumor” was a tumor-adjacent region on the same tissue block >1 mm from LUAD with a distinct invasive boundary separating off tumor regions from invasive LUAD.

### Immunohistochemistry

FFPE blocks were sectioned by microtome and slides deparaffinized and rehydrated. For RFP, Vimentin, NKX2-1, p-FAK, PCNA, and TNC staining, antigen retrieval was in Citrate buffer, pH 6.0. For TE-7, slides were incubated in EDTA, pH 8 retrieval buffer, and stained overnight, 4°C. Slides were developed with ImmPRESS HRP Horse anti-rabbit IgG Polymer Kit, ImmPACT DAB Substrate Kit, and Harris Hematoxylin counterstain. For sequential staining, coverslips were removed in xylene and slides re-stained using BOND Polymer Refine Red Detection kit using an automated Leica BOND slide stainer. anti-CD31 was on slides previously stained for TE7. anti-p-FAK on slides stained for TNC.

### Multiplex Immunofluorescence and analysis

Human tissue sections stained for TNC, p-FAK, and pan-keratin AE1/AE3 using the Leica Bond Biosystem. Anti-TNC was paired with the Opal 520 secondary, anti-p-FAK was with Opal 620, and AE1/AE3 with Opal 690. DAPI labeled the nuclei.

### Quantification of IHC using QuPath

For quantification of TNC in mouse tissue, off-tumor, hyperplasia, adenoma, and LUAD annotations were identified in tdTomato-stained slides, applied to the TNC-stained slides, and TNC was quantified by *Positive cell detection*. Off tumor regions were circular annotations with no transformed cells, ≥ 1,000,000 μm^2^. Hyperplasia were ≥20 connected transformed cells along alveolar walls. Adenoma and LUAD were as described under *Kras-driven Mouse Models*. The H-score, a normalized DAB intensity, was calculated in each annotation based on the cell mean optical density sum. 300 indicates 100% of cells exhibited the highest staining. For the tumor edge, LUADs were segmented into 60 µm “edge” annotations inside the tumor boundary using the expand annotation tool.

For quantification of extracellular TNC in human tissue, we trained a model to separate tumor, inflammation, and stroma and calculated TNC in the stroma. First, tumor boundaries and invasive regions were identified on H&E slides by the study pathologist (L.L.E.) and used as guides for manual annotation of five ROIs/case from off-tumor and invasive regions. H&E and TNC immunohistochemistry serial sections were co-registered and ROIs transferred to the TNC images. Next, a pixel-based random tree model excluded the cancer and immune cells from the TNC intensity calculation. The study pathologist annotated 10+ regions of cancer cell, immune cell, and stroma as the ground truth for model training and reviewed the classification output. The model was applied to the ROIs to exclude the immune and cancer regions. Cells in the stromal region were detected by *StarDist* nuclear segmentation. Mean staining intensity and total stromal area was obtained by the QuPath *add intensity features* function. Mean intensity of TNC = (mean staining intensity* total area of stroma) – (sum of mean cell staining intensity*cell area) / (total area of stroma – sum of cell areas).

For quantification of p-FAK in human tissue, the total tumor and 4 invasive areas of the tumor were marked on the H&E serial section by a board-certified lung pathologist and transferred to the fluorescent image. Four TNC-positive ROIs were automatically detected in the fluorescent image and expanded 20 μm. TNC-negative ROIs inside the tumor and similar in area to the positive regions were manually annotated. Cells were detected by the *StarDist* nuclear segmentation on the DAPI signal and classified as AE1/AE3 positive (epithelial) or negative. p-FAK H-score was quantified in extra-nuclear regions of the AE1/AE3 postive cells within the TNC-positive and -negative ROIs and intensity per cell was averaged.

For quantification of invasion based on tdTomato+ cells outside of the main tumor mass, tumors were annotated by an investigator blinded to their TNC status. A 150 μm-expanded ring around each tumor defined the tumor microenvironment. *Positive Cell Detection* within the tumor microenvironment quantified the percent tdTomato+ cells, which normalized the invading cell number to the area.

### In Situ Hybridization and quantification

RNAscope RED Assay with Immunohistochemistry Integrated Co-Detection (ACD) was following manufacturer’s protocol. Sections were baked at 60°C, deparaffinized, treated with H_2_O_2_, permeabilized with ACD co-detection antigen retrieval reagent at 100°C, incubated with the primary antibody overnight, 4°C, and treated with protease. Probes were incubated at 40°C, chemically amplified, labeled with alkaline phosphatase conversion of FastRED dye, and then slides incubated with IMPRESS HRP, DAB, and counterstained with Gills Hematoxylin I.

For manual counting, particles/cell of *TNC* were counted for each ROI. For human LUAD, 13 invasive and 12 off-tumor ROIs from 7 cases were quantified. Regions of invasive LUAD and off-tumor lung on the same slide were annotated by a lung pathologist. Tumor cells were TE7-with epithelial morphology and enlarged nuclei.

For automated counting in QuPath, the H&E, TNC, and TE7 images for each same were co-registered by the ImageCombinerWarpy extension to create a new QuPath entry with the combined registered images. Tumor areas were identified in the H&E image by the study pathologist and 5 regions of interest (ROIs, 1×1 mm) were manually selected in the H&E image by an independent investigator in an unbiased fasion. The outlines of the ROIs were transferred to the *TNC* ISH/IHC images. Within each ROI, we generated binary masks of fibroblasts (TE7+) and tumor and immune cells using the trained model described under *Quantification of IHC using QuPath*. Cell borders were obtained using nucleus segmentation and outline expansion by 5 µm. Subcellular spot detection identified the red *TNC* mRNA objects. Cells were classified into negative, 1+ *TNC* particles/cell, 4+ *TNC* particles/cell, and 10+ *TNC* particles/cell.

### Tumor burden and grade

Total lung and tumor areas were annotated in QuPath by pixel classification. Non-lung tissues including lymph nodes, large airways, and cell clusters on the outside surface of the lungs were excluded from analyses. Tumor burden was calculated as sum tumor area / sum lung area * 100. For tumor grade, H&E images were resampled to 0.5022 μm/pixel and processed through GLASS-AI using default settings. Entire lungs were analyzed for 10 week tumors, the largest lobe for 15 week tumors. Tumors were manually selected for analysis within the software to avoid artefacts with the identification of immune cells.

### Bioinformatics analysis of TNC expression

For TCGA samples, RNA-seq expression data for 505 human tumors was downloaded from the GDC data portal (TCGA-LUAD) using TCGAbiolinks. Tumors missing stage or technical replicates or FFPE were removed. mRNAs with 10 or fewer raw counts in 95% of samples were removed. For the remaining 17,275 proteins, fold change of mRNA counts was estimated using DESeq2. Differential expression was tested with a negative binomial generalized linear model, 5% false discovery rate. All human tumors reported 3 or more counts for *TNC*. Tumor purity and stromal content were calculated for the 227 annotated TCGA samples using ESTIMATE.

For GSEA, TCGA-LUAD samples from the 505 human tumors were split into quartiles. Differentially expressed genes between the TNC high and TNC low quartiles were compared to reported gene sets. Genes from each signature were mapped to Ensembl. GSEA calculations were in R using the fast GSEA (fgsea) algorithm, filtered using a 10% FDR, and plotted according to GSEA score. Leading edge genes in the TNC signature were processed in ShinyGO 0.80 using the GO Biological Process Pathway database.

For KP tumors, scRNAseq data from labeled KP tumors^41^ was downloaded with sample and cell barcodes and cluster names and processed using Seurat’s *NormalizeData* algorithm and the log normalized counts/cell plotted.

For *TNC* in human lungs, scRNAseq data from Sikkema et al.^42^ was analyzed using the online tool at CZ CellxGene and by extracting data subsets using Scanpy from the downloadable .h5ad file for graphing in R. Plotting of cell types was limited to ≤1000 cells.

### TNC isoforms

scRNAseq Fastq files from 4 treatment naive primary human patient LUAD samples with greater than 10 cells from Maynard, et. al^48^ were demultiplexed by cell barcode (LT-S52, LT-S56, LT-S67, and LT-S74). Files were aligned to Hg38 using a *Next-Flow* pipeline with *STAR* alignment using the *samtools* module to prepare BAM files of aligned sequences. Expression of individual *TNC* isoforms was compared between tumor cells and fibroblasts, as previously annotated^48^, by RSEM using *rsem*.

### Cell lines and culture

Cells were maintained in DMEM supplemented with 10% FBS, at 37°C, 5% CO_2_ and periodically tested negative for mycoplasma. Human *RAS* mutant LUAD cell lines were H1299, A549, and H23. Mouse LUAD cells were 1783, 3658, 4043, 7865, from Kras^G12D^;Trp53^Null^; Nkx2-1^Null^ tumors. Normal human lung fibroblasts (NHLF) and cancer-associated lung fibroblasts (CAF) were cultured in Lonza Fibroblast Media and MSCGro media, respectively and after two passages, immortalized by infection with retrovirus expressing pBabe-hTERT-p53^DD^ generated in Phoenix-AMPHO cells, then cultured in DMEM, 5% FBS. Mouse embryonic fibroblasts: clonetech MEFs and m28 from C57BL6 mice. Plate coatings were BSA (1% in PBS) and TNC (10 μg/μl). Treatments with inhibitors or blocking antibodies were 3 hours prior to assay.

### Immunoblotting

Cells were lysed in RIPA with Halt protease and phosphatase inhibitor cocktail. Protein was normalized using Bradford Protein Assay Reagent, separated by SDS-PAGE gel, transferred to 0.45 μm Nitrocellulose, and probed with antibodies against TNC and Vinculin, followed by IRDye-conjugated secondary antibodies. Westerns were visualized by Odyssey CLx Imaging System (LI-COR) and quantified in Image Studio.

### p-FAK Immunofluorescence

Cells were fixed in 4% Formaldehyde in PHEM buffer (60 mM PIPES, 25 mM HEPES, 10 mM EGTA, 4 mM MgSO_4_, 50 mM β-glycerophosphate, 0.2 mM Vanadate, pH 6.9), permeabilized in 1% CHAPS in PHEM, blocked in MBST (50 mM MOPS, 150 mM NaCl, 0.05% Tween-20, pH 7.4) with goat serum, and incubated with primary and secondary antibodies in MBST with goat serum. Images were on a Nikon Ti inverted microscope with a Plan Fluor ELWD 20× air objective with Andor Zyla cMOS camera using Nikon Elements. Fluorescence intensity was measured in FIJI as IntDen.

### Cell proliferation and migration

Proliferation was with Janus Green B with three technical replicates read on an Epoch2 (Biotek) plate reader at 620 nm.

For migration, cells were cultured on glass-bottomed 12-well plates in FluoroBrite DMEM with 10% FBS, 20 mm HEPES, and DRAQ5 and imaged every 10 mins for 15 hours on a Nikon Ti inverted microscope with a Plan Fluor ELWD 20× air objective described above, with an Okolabs environmental chamber. Cells were tracked using the DRAQ5 and custom software based off of u-track multiple-particle tracking. Migration velocity was calculated as the Mean Squared Displacement (MSD), the average square displacement over increasing time intervals. Cells with persistent random walk motion, in which the MSD increases in a superdiffusive manner (MSD(t) ∝ t^α^, where 1<α<2) were included.

### Atomic Force Microscopy

Lungs were inflated with PBS/O.C.T., embedded in O.C.T., and frozen in 2-methylbutane. 10 μm sections were cut using a Leica CM1860 UV cryostat and mounted on glass slides. Areas of interest were identified under phase contrast and fluorescent microscope with at 40× and 200× objectives. All tumors analyzed were approximately the same size. AFM measurements were on rehydrated tissue slices using a Catalyst Bioscope atomic force microscope (Bruker) and the MIRO 2.0 extension through Nanoscope 9.1 software, with borosilicate sphere AFM tips with 2.5 μm radius (Novascan) and spring constant estimated at ∼100 pN/nm by thermal tune. Force curves were performed randomly in 150 x 150 µm² areas. Elastic modulus (Young’s modulus) was estimated by fitting force curves (NanoScope Analysis 2.0 software, Bruker) with the Hertz contact model: 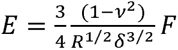 with R the tip radius, ν the Poisson’s ratio assumed at 0.4 for lung tissue^74^ and δ the sample indentation. For each sample, three areas of interest were analyzed from non-consecutive tissue slices. For each area, 25 force curves were randomly performed, analyzed to determine their elastic values, and averaged to report one elastic value.

### PCLS Immunofluorescence and analysis

Mice were euthanized by CO_2_ and lungs inflated with warm 1.5% low melt agarose in PBS, administered through the trachea. Lung tissue was harvested in culture media (DMEM/F12 without phenol red, supplemented with 0.1% FBS, 0.1 μg/ml Hydrocortisone, 0.1μg/ml EGF, and 1x Pen-Strep) and sliced on a vibratome. PCLSs were fixed in 4% formaldehyde, permeabilized, stained with CNA35-eGFP, anti-TNC, or HUTS-4, and imaged on a Leica Sp8 white light laser confocal microscope using an HC PL APO 10×/0.40 objective or an HC PL APO CS2 40x/1.10 water objective with immersol for collagen analysis. Image acquisition was 1024×1024 resolution using 1.14 μm pixel size, 1-2 μm z-step, excitation at 488 nm, 554 nm, and 653 nm, and emission 493-550 and 564-658 and 658-778. Image stacks were visualized in Fluorender.

For collagen analyses, a square confocal voxel was obtained with 0.3 μm pixel size and 0.3 μm z-step. In MATLAB, binary masks were calculated for each 2D z-slice of the image volume. Mask skeletonization yielded the collagen fiber centers and the distance transform of the skeletonization yielded the fiber radius, which was doubled to obtain the 2D collagen thickness. Collagen pore size was calculated by creating 2D binary masks for pores with x-y plane cross-sections <100 μm^2^ in each z-slice and building 3D pore binary masks by iteratively combining 2D masks in the adjacent z-slices. 3D masks were rendered into 3D triangulated meshes. The Dijkstra algorithm identified the 3D mask centerline, which was used to find the pore radius from the distance transform.

Quantification of HUTS-4 and TNC was in MATLAB as an image volume, resampled to a 1 μm^3^ voxel. TNC and tdTomato-positive tumor cells were masked. The TNC mask was expanded 2 μm to create a shell areas of extracellular TNC for analysis. The tumor cell masks were expanded concentrically outward by 0.5 μm to capture the HUTS-4 intensity at the cell surface. The sum intensities for TNC and HUTS-4 in the overlapping voxels in the TNC-adjacent shell and expanded tumor surface were plotted.

### Tumor recurrence

Patient clinical data were collected by affiliates of the Oncology Research Information Exchange Network (ORIEN, https://orien.tcc.org) using harmonized data fields within a RedCap database. Clinical data was matched to sequenced sample data and patient identifiers using a custom-built Python 3.9 script. Treatment agnostic progression free survival was calculated from the patient’s date of surgery to progression events based on documented progression or recurrence, with patients censored at the age of last contact. Progression and recurrence included clinical notes of lesion growth, medication changes due to progression, and metastasis specifically within the lung or pleura. Early progression events were filtered to exclude events before 45 days to account for any lag in response or early interventions and exclude events involving metastasis. All clinical and molecular data were loaded into an institutional instance of cBioPortal version 5.4.2 for review and analysis.

For the 547 samples with RNAseq data for *TNC*, the *TNC* raw counts were normalized using DESeq2 and added to the assembly in cBioportal. The *TNC*-sequenced samples were further filtered to exclude those noted to be stage III or IV, collected from connective or subcutaneous tissue, with death due to causes other than cancer, or neuroendocrine, squamous, basaloid squamous, or adenosquamous histology. Samples were also filtered to include only those with 30-90% tumor content to ensure inclusion of both the tumor and tumor microenvironment and those with 10-120 months (10 years) since the last follow up to ensure sufficient time for recurrence. Cases with metastasis outside of the lung, lung lymph nodes, pleura, or thorax were excluded. Cases were divided into those with the lower 25% and upper 75% of TNC expression (≤12.85 or > 12.85 log2 normalized counts) and plotted as a survival curve for time to first progression with log-rank test for significance.

### Plotting and Statistics

Plots and statistics were in GraphPad PRISM or MATLAB. In distributions, the central line marks the median. Notches in box plots show 95% confidence interval. Normality was tested with the Shapiro-Wilk test. For two independent groups with normally distributed data, a two-tailed T-test with Welch’s correction for unequal variance tested significance. For multiple groups with normal data, significance was by one-way ANOVA with Tukey’s posthoc test. For data with deviations in normality, two independent groups were compared with the non-parametric Mann Whitney test or multiple groups with the Kruskal-Wallis one-way ANOVA with Sidak’s posthoc test. Paired data was tested with the two-sample non-parametric Wilcoxon matched-pairs signed rank. Distributions with unlimited sample size and where extreme values confer biological phenotype were tested with the two-sample nonparametric K-S test. *p<0.05, **p<0.01, ***p<0.001, and ****p<0.0001. Bioinformatics analyses in DESeq2 used the Wald test with the Benjamin and Hochburg method of multiple testing for adjusted p values.

## Materials

Key resources table:

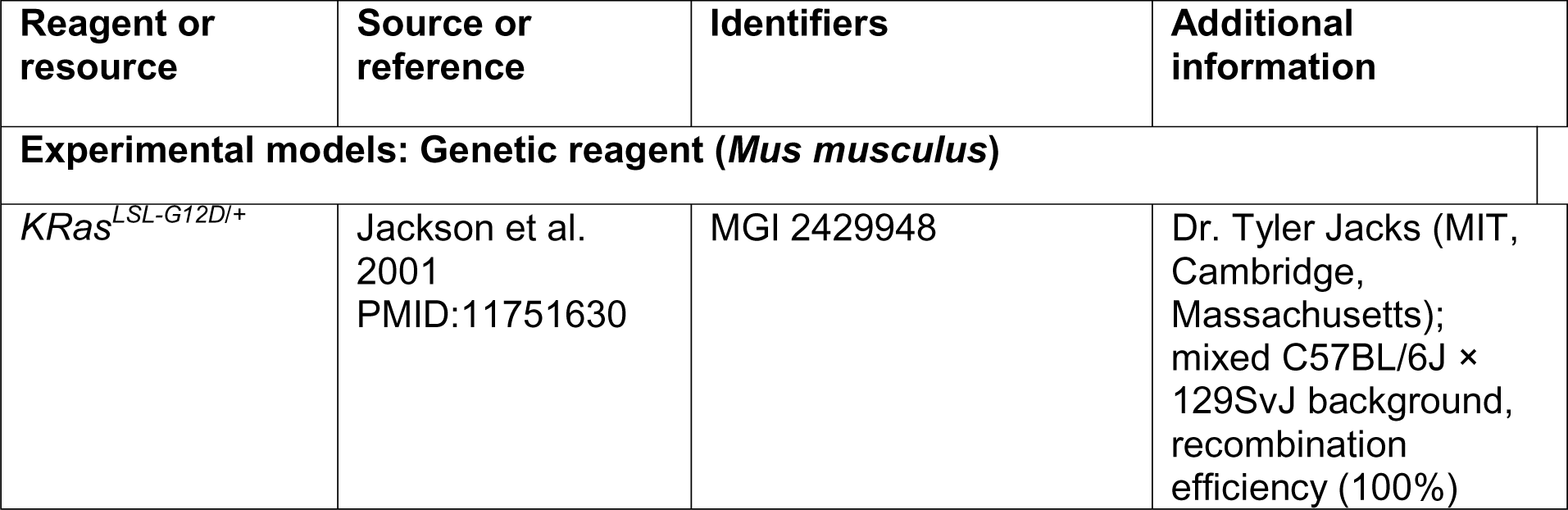
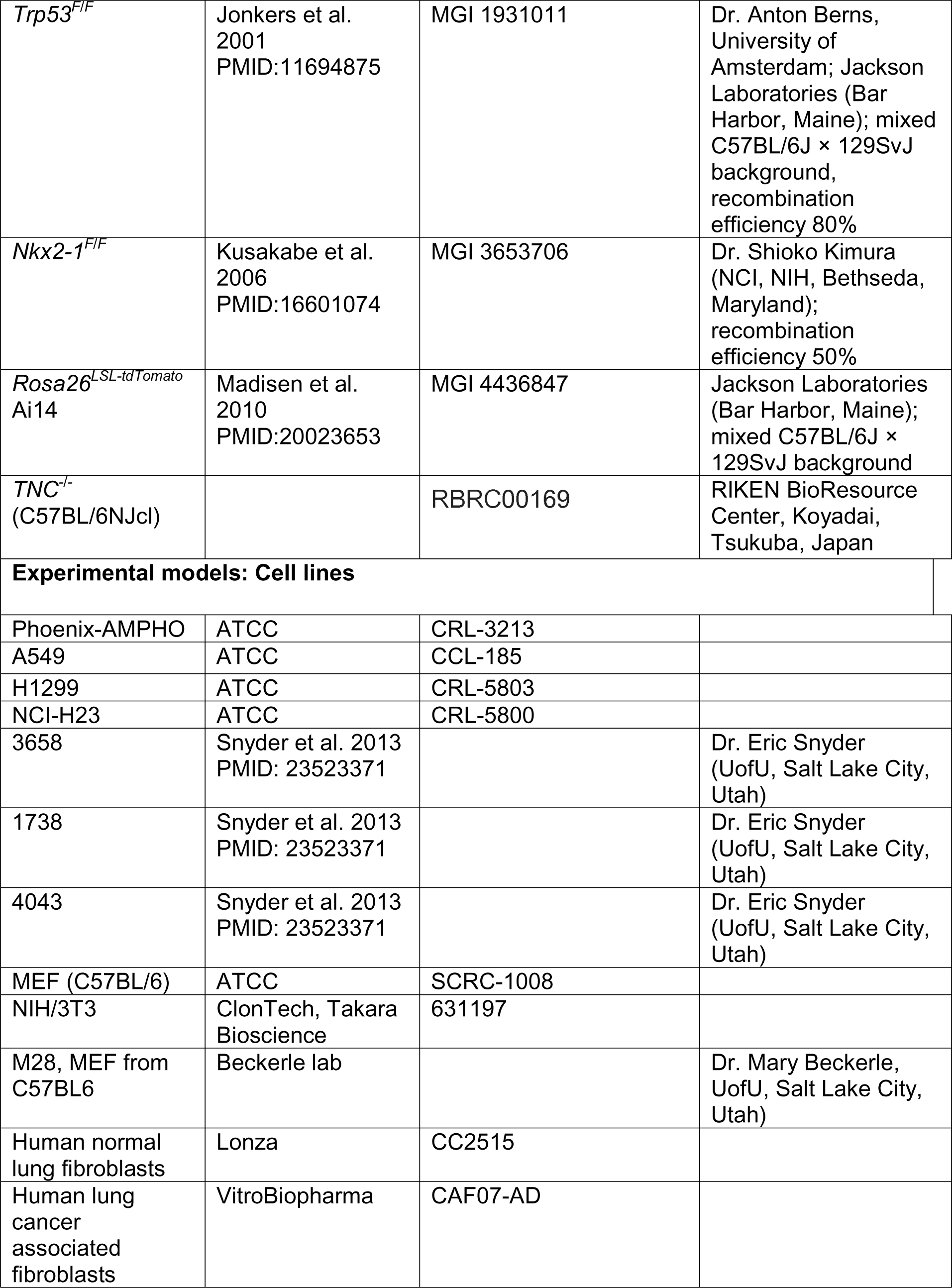
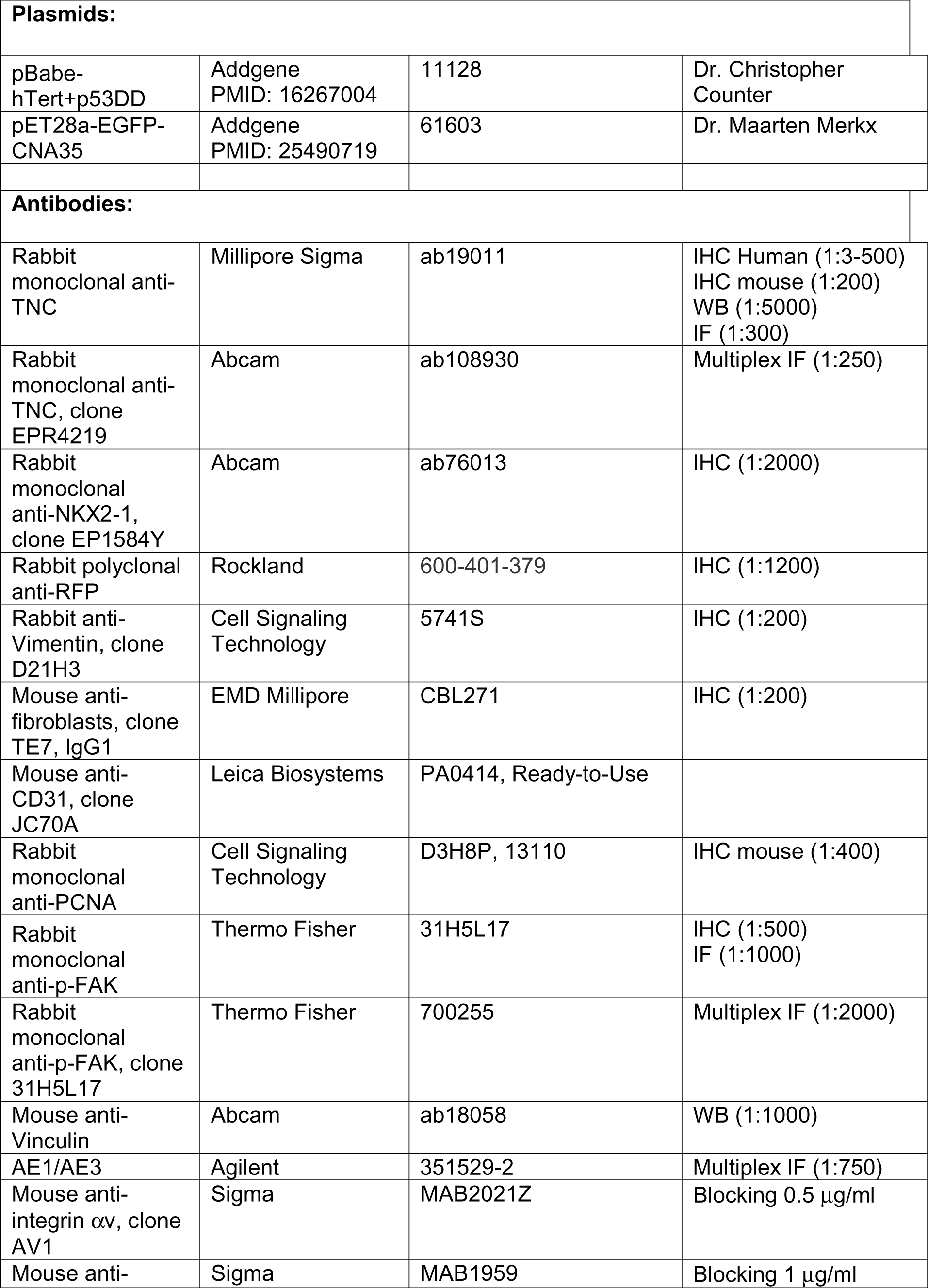
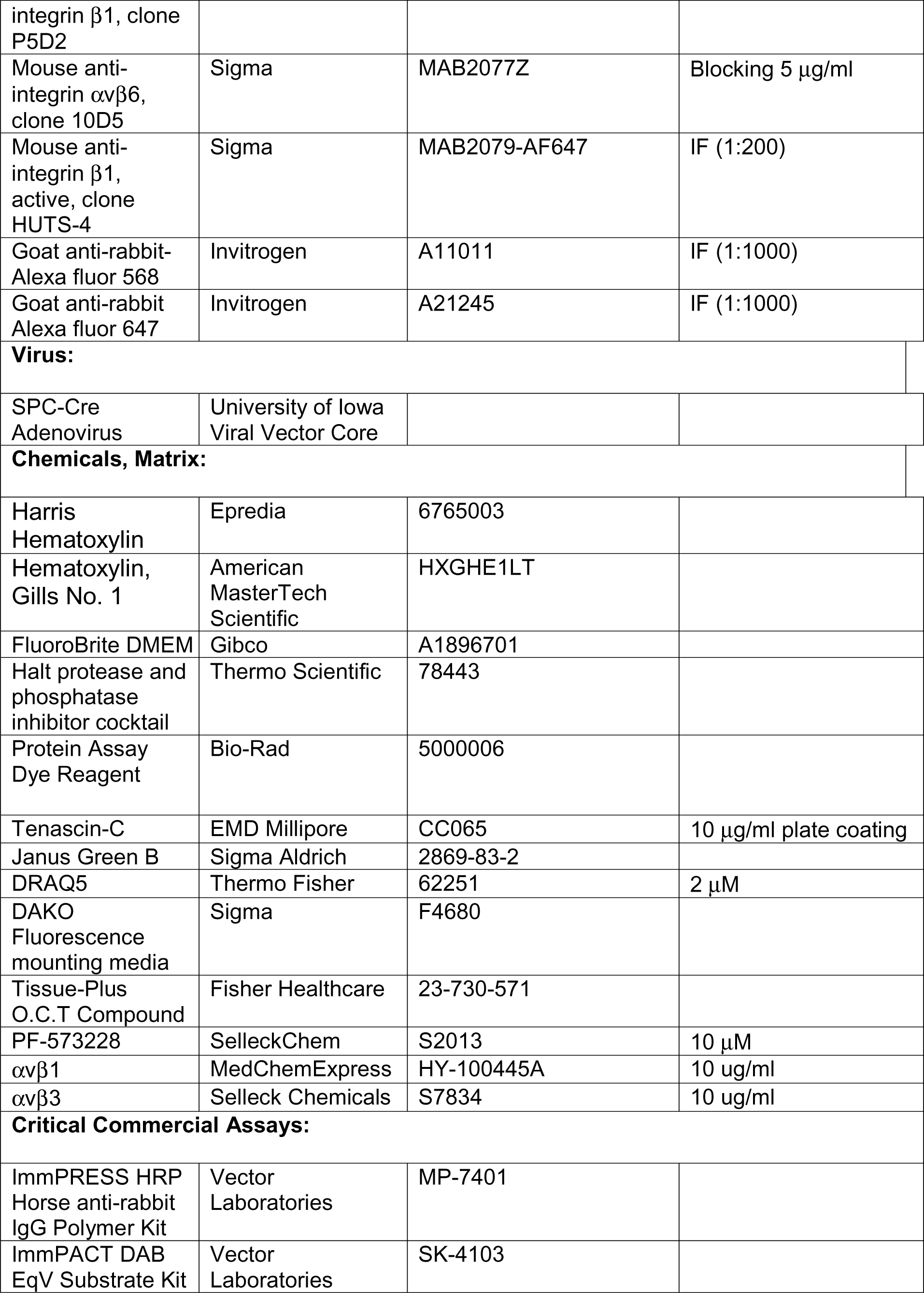
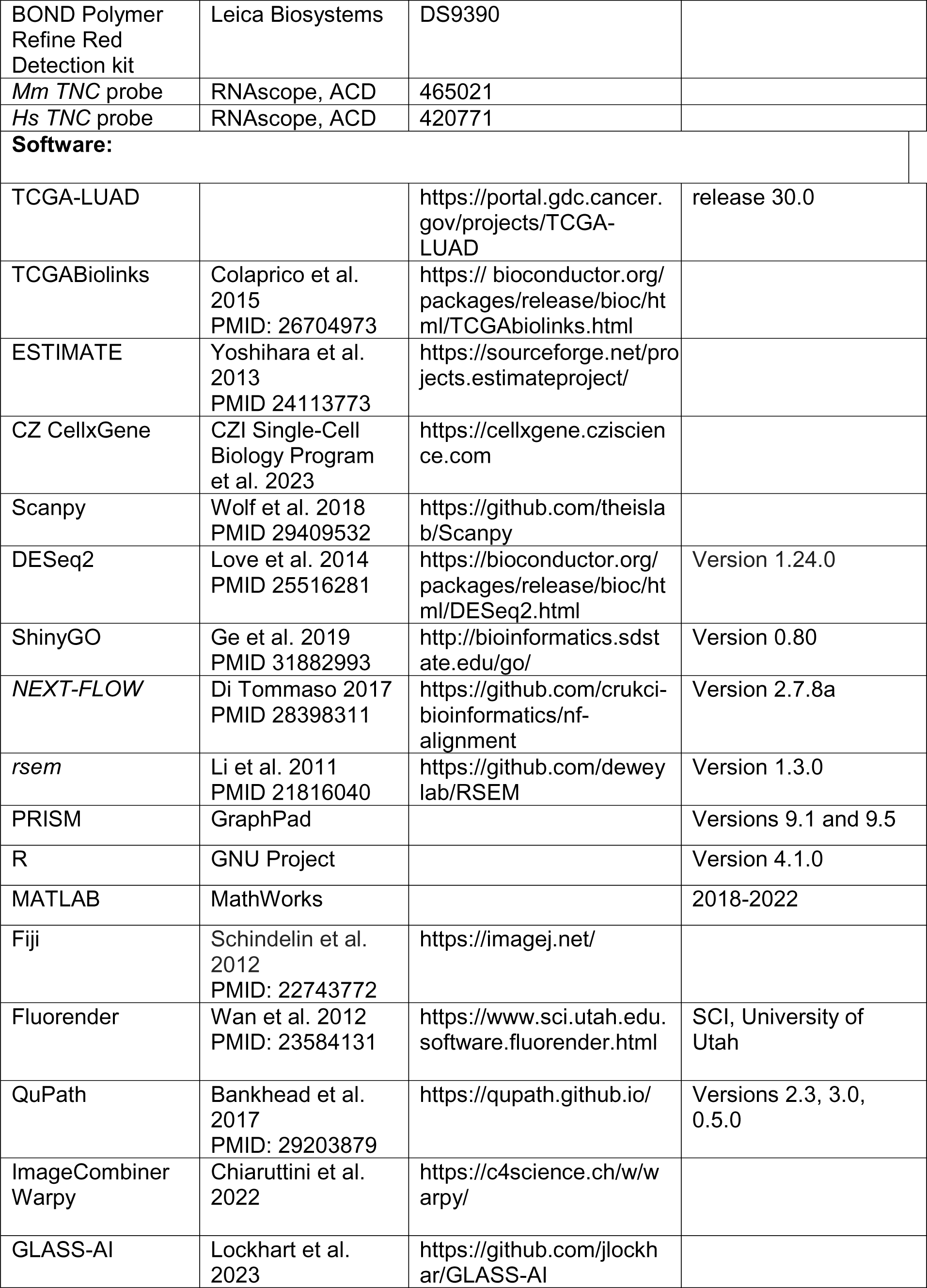
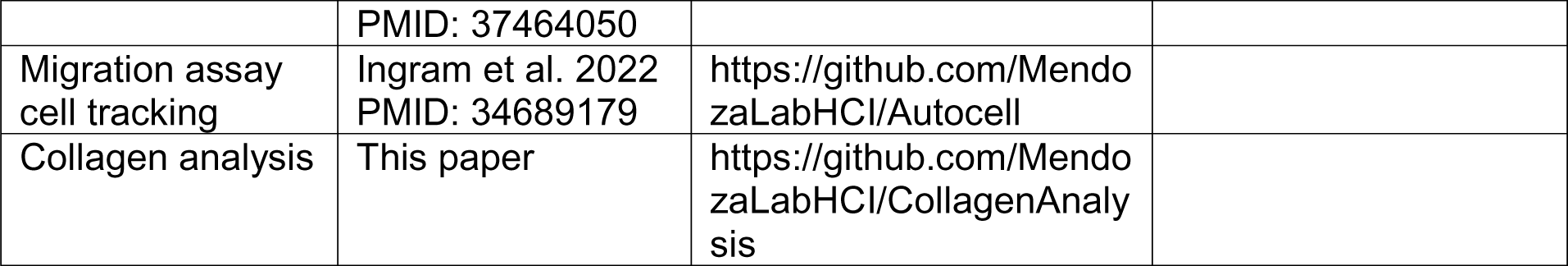

